# A foundation model for bioactivity prediction using pairwise meta-learning

**DOI:** 10.1101/2023.10.30.564861

**Authors:** Bin Feng, Zequn Liu, Nanlan Huang, Zhiping Xiao, Haomiao Zhang, Srbuhi Mirzoyan, Hanwen Xu, Jiaran Hao, Yinghui Xu, Ming Zhang, Sheng Wang

## Abstract

Compound bioactivity plays an important role in different stages of drug development and discovery. Existing machine learning approaches have poor generalization ability in compound bioactivity prediction due to the small number of compounds in each assay and incompatible measurements among assays. Here, we propose ActFound, a foundation model for bioactivity prediction trained on 2.3 million experimentally-measured bioactivity compounds and 50, 869 assays from ChEMBL and BindingDB. The key idea of ActFound is to employ pairwise learning to learn the relative value differences between two compounds within the same assay to circumvent the incompatibility among assays. ActFound further exploits meta-learning to jointly optimize the model from all assays. On six real-world bioactivity datasets, ActFound demonstrates accurate in-domain prediction and strong generalization across datasets, assay types, and molecular scaffolds. We also demonstrated that ActFound can be used as an accurate alternative to the leading computational chemistry software FEP+(OPLS4) by achieving comparable performance when only using a few data points for fine-tuning. The promising results of ActFound indicate that ActFound can be an effective foundation model for a wide range of tasks in compound bioactivity prediction, paving the path for machine learning-based drug development and discovery.

## 1 Introduction

Compound bioactivity plays an important role in drug development and discovery due to its broad applicability in different stages of drug discovery, including hit identification,^1–3^ drug repurposing,^4^ and lead optimization.^5, 6^ Bioactivity prediction aims to predict the activity of compounds in a specific binding type, functional type, or pharmacokinetic type assay,^7^ allowing scientists to rapidly identify desired compounds from a large number of samples.^8^ Traditional computational chemistry approaches rely on 3D target-protein structure^9^ to predict bioactivity and suffer from the inherent trade-off between accuracy and computational cost.^10^ For example, state-of-the-art methods, such as free energy perturbation (FEP),^11^ can provide accurate predictions for several binding type assays,^12^ but also require extensive computational resources that are often not affordable for large-scale applications.^9^

In the past decade, deep learning methods have demonstrated great potential in bioactivity prediction by using fewer computational resources compared to computational chemistry approaches.^13^ However, existing deep learning approaches have two limitations. First, only a few compounds (12.7 compounds in ChEMBL^7^ and 54.3 compounds in BindingDB^14^ on average) are measured in each assay due to the expensive experimental cost. As a result, there is not enough training data to train an assay-specific model. Second, although advanced machine learning techniques, such as transfer learning,^15–20^ multi-task learning^21^ and meta-learning,^22–28^ can be used to jointly learn from all assays, these assays are often incompatible in terms of units (e.g., nMol, mg/kg/day, ug/ml), value ranges, and measurements (e.g., IC50, DC50). This leads to incorrect transfer between assays and hurts the prediction performance, especially for rare compounds and assay types.

Recently, foundation models have achieved promising results in computer vision and natural language processing.^29–31^ Foundation models aim to learn a pre-trained model from large-scale and diverse datasets that serve as a foundation for a variety of downstream tasks. The key advantage of foundation models is that they are capable of few-shot learning, which means that they only need a few labeled data to be fine-tuned for downstream tasks.^32, 33^ Because of this label-efficient advantage, foundation models have also been applied to biomedicine, where labeled data are often expensive.^34^ We hypothesize that a foundation model could also be effective for bioactivity prediction because it presents large and diverse assays while each assay only contains a few compounds.

In this work, we propose ActFound, a foundation model for bioactivity prediction. Our foundation model is developed based on two machine learning techniques: meta-learning and pairwise learning. Meta-learning is employed to jointly train the model from a large number of diverse assays, making it an initialization for new assays with limited data.^22–24^ Pairwise learning is used to address the inherent incompatibility among assays. Our intuition is that although compounds from different assays may have varying units, value ranges, or measurement metrics, those within the same assay are comparable. Instead of directly predicting absolute bioactivity values, we propose to learn the relative bioactivity difference between two compounds within the same assay using pairwise learning.^35, 36^ Specifically, for each assay, we utilize a Siamese Network architecture to acquire the relative difference in bioactivity values between two compounds. This approach allows us to reconstruct the bioactivity values for previously unseen compounds using the predicted relative bioactivities.

We evaluated ActFound on four downstream bioactivity prediction tasks: in-domain bioactivity prediction, cross-domain bioactivity prediction, FEP prediction, and cancer drug response prediction. On in-domain bioactivity prediction, ActFound outperformed all nine competing methods on ChEMBL, BindingDB, FS-Mol, and pQSAR-ChEMBL. On cross-domain bioactivity prediction, where the model is trained in one domain (e.g., ChEMBL) and tested on another domain (e.g., BindingDB), our model also achieved the best results, indicating that ActFound is agnostic to measurement units and can be applied to different types of assays, and we also provided practical guidance on when to use ActFound for researchers. Furthermore, we demonstrated that ActFound can be an alternative to FEP calculation, a resource-intensive Relative Binding Free Energy (RBFE) computation chemistry method. Specifically, we found that by using 40% of the compounds for fine-tuning (12 compounds on average for each assay), our model can surpass FEP+(OPLS4),^37^ the leading commercial software for FEP calculation, by 9.5% in terms of RMSE on FEP benchmarks. Lastly, we showed that a cancer drug response regressor pre-trained using ActFound exhibited promising performance on zero-shot drug sensitivity prediction, underscoring the broad applicability of ActFound. Collectively, ActFound stands as a foundational model for bioactivity prediction developed through pairwise meta-learning, and can be broadly applied to a variety of applications in drug development and discovery.

## 2 Results

### 2.1 Overview of ActFound

ActFound is a foundation model for bioactivity prediction. To effectively utilize a large number of available assays and compounds, ActFound is developed to address two major challenges in bioactivity prediction, the limited availability of labeled data for each assay and the incompatibility among different assays. With only a few compounds and their corresponding experimental bioactivity on a new assay, ActFound can accurately predict the bioactivity of new compounds in the assay. ActFound provides a pre-trained bioactivity model using meta-learning and pairwise learning (**Fig. 1a**). At the pre-training stage, diverse assays are used as “meta-training” datasets. The parameters of ActFound are interactively updated on these assays using a bi-level optimization process, which consists of an inner loop for fine-tuning specific assays, and an outer loop for updating model parameters based on the performance of fine-tuned models. As a result, ActFound is more sensitive to the updates from various assays and can later quickly adapt to new assays. The backbone of ActFound is a Siamese Network calculating the relative difference in bioactivity values between two compounds, allowing us to circumvent the issue of incompatible bioactivity values across different assays and focus on the compatible relative activity. The final bioactivity values can be reconstructed based on the predicted relative bioactivity values (**Fig. 1b**). At the fine-tuning stage, ActFound is fine-tuned using a few compounds with experimental bioactivities in the new assay, and then predicts bioactivity predictions on unmeasured compounds (**Fig. 1c**). We demonstrated that ActFound can accurately perform in-domain bioactivity prediction and exhibit strong generalization ability across domains.

**Figure 1.**
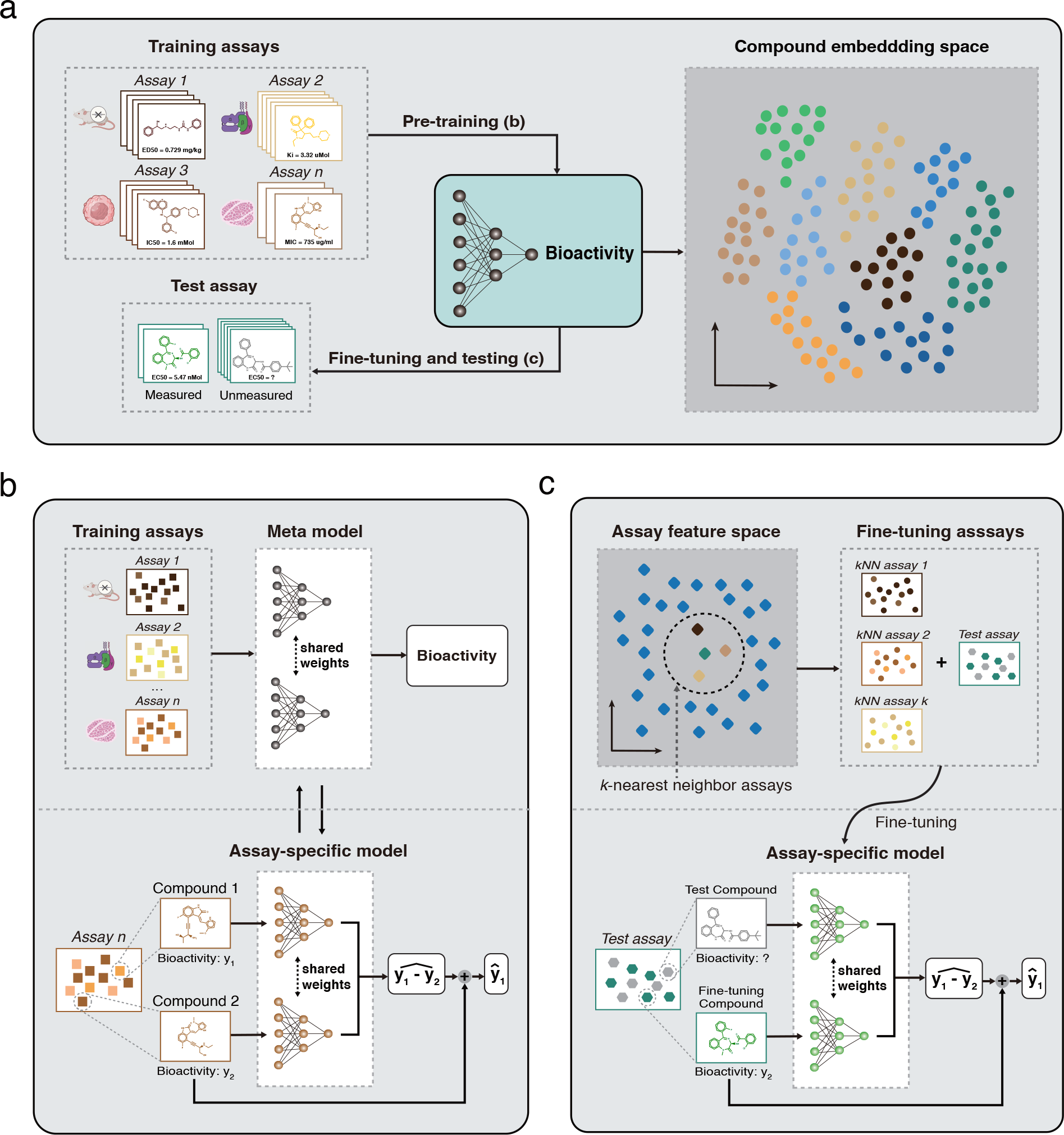
Overview of ActFound. **a**, ActFound is a foundation model for bioactivity prediction pre-trained on a large number of assays. After pre-training, ActFound can be fine-tuned on a test assay with just a few measured compounds, and then predict bioactivity values for other unmeasured compounds in this assay. The pre-training stage enables ActFound to project the compounds from diverse assays to a shared embedding space. **b**, ActFound exploits pairwise meta-learning for pre-training strategy. A Siamese Network is used to learn the relative bioactivity value differences between two compounds to address the incompatibility among assays. The final bioactivity value can be calculated using the predicted relative bioactivity value difference. **c**, ActFound employs a *k*-nearest neighbor-based fine-tuning strategy where the *k*-nearest assays are identified to jointly fine-tune the test assay. This makes the pre-trained model quickly adapt to the test assay, improving the generalizability of ActFound.

### 2.2 Accurate in-domain bioactivity prediction

We first sought to evaluate the performance of ActFound for bioactivity prediction in the in-domain setting (see **Methods**) on ChEMBL^7^ and BindingDB.^14^ Following previous work,^22^ we evaluated on the 16-shot setting, where 16 compounds in each assay were used to fine-tune the model, and the remaining ones were held out for evaluation. We found that ActFound outperformed all competing methods on both datasets in terms of *r*^2^ (**Fig. 2a**, *p*-value < 10^−11^) and RMSE (**Fig. 2b**, *p*-value < 10^−20^). First, ActFound substantially outperformed conventional meta-learning approaches MAML^38^ and ProtoNet,^39^ indicating the effectiveness of using pairwise learning to learn the relative bioactivity values. Second, we found that ActFound is better than ActFound(transfer), a transfer learning based variant of our method, demonstrating the advantage of using meta-learning to train a foundation model from diverse assays. Although being worse than ActFound, MAML and ProtoNet are better than other non-meta-learning approaches, again indicating the effectiveness of using meta-learning for this task. We found that the improvement of meta-learning-based approaches is larger on ChEMBL than BindingDB. Since ChEMBL has 35, 644 assays, more than twice the amount of assays compared to the 15, 225 assays on BindingDB, it can better exploit the advantage of meta-learning to quickly adapt the model to new assays, reassuring the prominence of using meta-learning to build a foundation model for bioactivity prediction.

**Figure 2.**
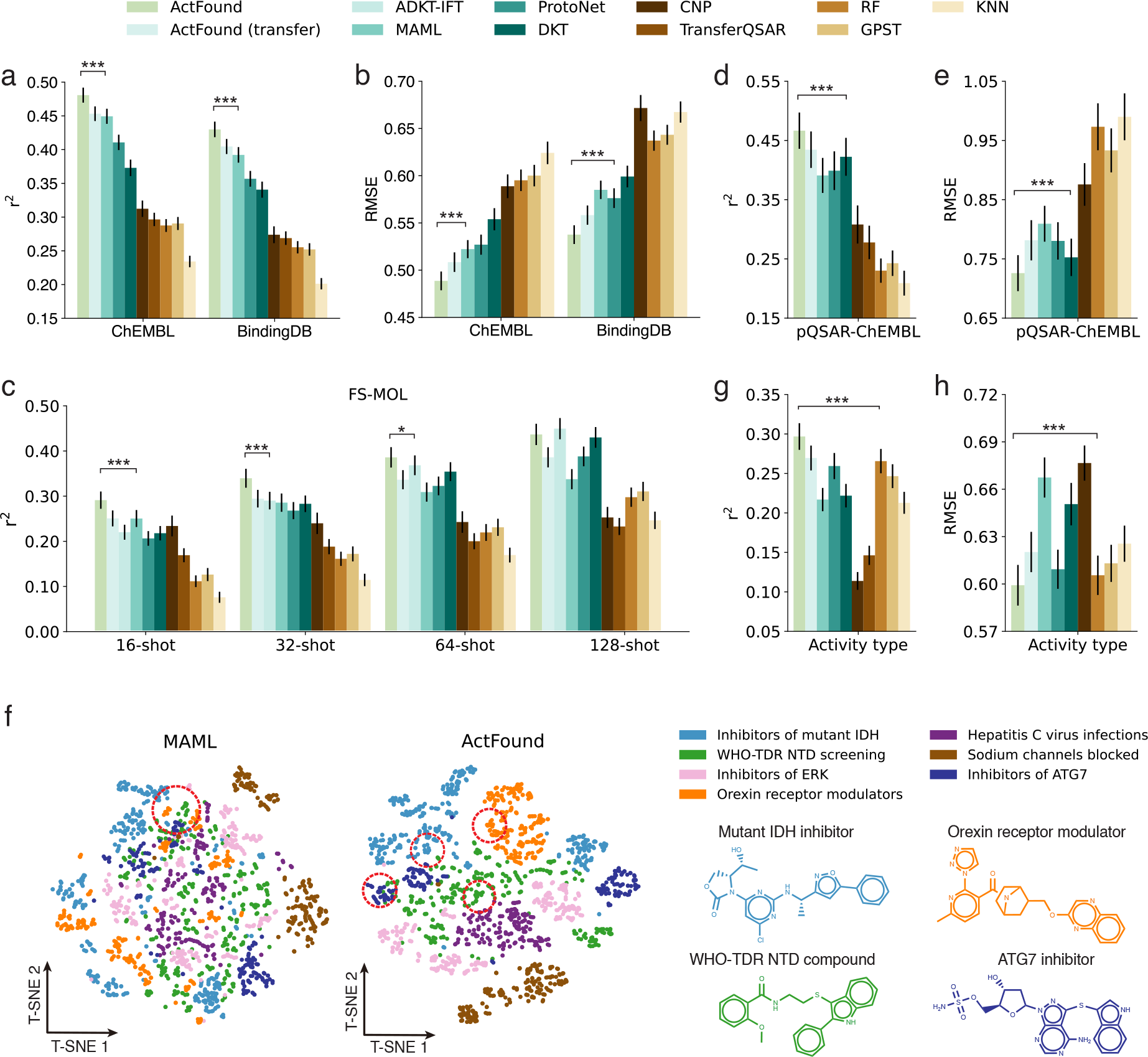
Evaluation of the in-domain bioactivity prediction. **a,b**, Bar plots comparing ActFound and competing methods in 16-shot bioactivity prediction on ChEMBL and BindingDB in terms of *r*^2^ (**a**) and RMSE (**b**). The asterisks indicate that ActFound outperforms the next best-performing model in the metric, with t-test significance levels of p-value < 5 × 10^−2^ for *, p-value < 5 × 10^−3^ for **, and p-value < 5 × 10^−4^ for ***. **c**, Bar plot comparing the 16, 32, 64, 128-shot bioactivity prediction on FS-MOL using *r*^2^. **d,e**, Bar plots comparing bioactivity prediction using scaffold data splitting on pQSAR-ChEMBL in terms of *r*^2^ (**d**) and RMSE (**e**), 75% of the experimental bioactivity is used for fine-tuning. **f**, T-SNE plots comparing compound embeddings obtained by ActFound (left) and MAML (middle). All compounds are from ChEMBL test set assays. Points are colored according to the assay, and assays with the largest number of compounds are selected. Four compounds which are denoted using red circles are illustrated (right). **g,h**, Bar plots comparing the bioactivity prediction on ChEMBL-Activity assays (unit=%) in terms of *r*^2^ (**g**) and RMSE (**h**).

Next, we evaluated our method FS-Mol.^22^ FS-Mol is curated from ChEMBL by only retaining assays with molar concentration units (e.g., nMol), thus excluding many costly *in vivo* assays measured in dosing units (e.g., mg/kg/day). FS-Mol is substantially smaller and only contains 13.8% of assays and 30.8% of unique compounds compared to ChEMBL. We assessed the performance using different number of fine-tuning compounds from 16 to 128 (**Fig. 2c, Supplementary Figure 1,2**) and found that ActFound achieved the best performance in the low-resource setting (16-shot, 32-shot) and comparable performance on 128-shot (*p*-value > 0.95). We attributed the improvement of our method to meta-learning, which is known to perform better when only limited labeled data are available. We further examined a more challenging scaffold data splitting setting on pQSAR-ChEMBL,^21^ where the fine-tuning and the test compounds of an assay do not have any overlapping molecular scaffolds. Since the molecular scaffold is the core structure that different functional groups attach to, this setting can more rigorously evaluate the generalizability of our method. ActFound again performs the best in this setting (**Fig. 2d,e, Supplementary Figure 3**), demonstrating its generalizability to unseen molecular scaffolds.

To better understand the superior performance of learning the relative bioactivity difference, we contrasted compound embeddings obtained by ActFound and MAML in **Fig. 2f**. Since most assays are developed for lead optimization, compounds in the same assay should have similar structures and thus be clustered together. Compared to MAML, embeddings from ActFound presented more visible patterns, confirming the high-quality embeddings learned from ActFound. Among all assays, we found that the performance of ActFound is the least desirable on assays that have the percentage (%) unit, whereas the training assays are molar concentration unit, density unit (e.g., ug/mL) or dosing unit (**Supplementary Figure 4**). Nevertheless, ActFound still surpassed other transfer learning and meta-learning approaches (**Fig. 2g,h**), again suggesting that our approach possesses better generalization capability across units. Collectively, the prominent performance of ActFound in the in-domain setting demonstrates the effectiveness of using pairwise meta-learning to build a foundation model for bioactivity prediction, further motivating us to investigate its performance in the more challenging cross-domain setting.

### 2.3 Improved generalization across domains

A key advantage of foundation models is the ability to generalize the prediction to new domains and datasets. After observing ActFound’s strong generalizable performance on unseen molecular scaffolds and units, we next systematically evaluated its generalization in the cross-domain setting, where the model is trained using all assays from one domain (e.g., ChEMBL) and fine-tuned on all assays from another domain (e.g., BindingDB). The results are summarized in **Fig. 3**. We first noticed that ActFound yielded a slight performance degradation in the cross-domain setting compared to the in-domain setting, demonstrating that this is a more challenging setting. Nevertheless, ActFound still outperformed all comparison approaches under both metrics on both datasets. For example, ActFound obtained 0.33 *r*^2^, which is 18% higher than the best-competing method MAML on the cross-domain prediction between ChEMBL and BindingDB (**Fig. 3a,b**). We found that the performance of Act-Found is better on ChEMBL-to-BindingDB than BindingDB-to-ChEMBL in terms of performance drops compared to the in-domain setting. Similar to our observation in the in-domain setting, we again attributed this to the larger number of assays in ChEMBL compared to BindingDB, which provides more tasks for our pairwise meta-learning framework.

**Figure 3.**
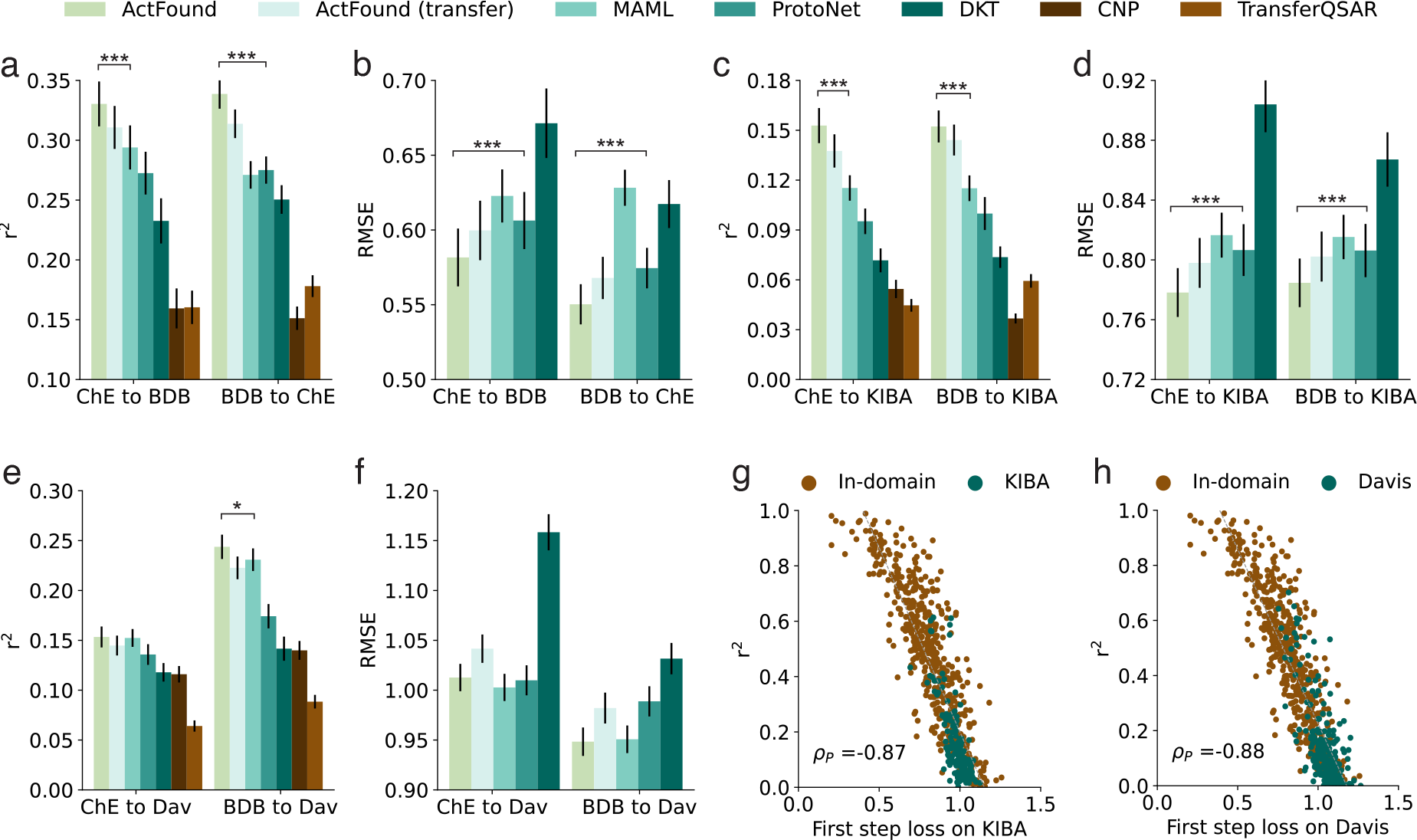
Evaluation of the cross-domain bioactivity prediction. **a,b**, Bar plots comparing the cross-domain bioactivity prediction between ChEMBL and BindingDB in terms of *r*^2^ (**a**) and RMSE (**b**). ChE, BDB are abbreviations for ChEMBL and BindingDB respectively. **c-f**, Bar plots comparing the bioactivity prediction on KIBA and Davis in terms of *r*^2^ and RMSE, and Dav is an abbreviation of Davis. **g,h**, Scatter plots comparing the loss for the first step and the test performance in terms of *r*^2^ on KIBA (**g**) and Davis (**h**).

Since BindingDB and ChEMBL are both collected from diverse sources, there might exist overlapping assays that cause data leakage. In addition to our efforts to exclude these related assays (see **Methods**), we exploited two independent kinase inhibitor datasets KIBA^40^ and Davis^41^ to eval-uate the cross-domain prediction performance. We first noticed that the performance on these two datasets is substantially worse than the performance on BindingDB and ChEMBL, suggesting them to be more challenging tasks. Our method achieved the overall best performance on both datasets (**Fig. 3c-f, Supplementary Figure 5,6**), reassuring the superiority of pairwise learning and meta-learning. Moreover, the performance enhancement of ActFound relative to MAML is larger on KIBA than Davis. In contrast to David, measurement units in KIBA do not exist in ChEMBL or BindingDB, again demonstrating that ActFound is agnostic to measurement units and can be better generalized to assays with never-before-seen measurement units.

Because the performance of our method and competing approaches all dropped in the cross-domain setting compared to the in-domain setting, it is beneficial to let end users know which test assays our method can perform well. We found that the test performance of the test assay is strongly correlated with the loss value for the first optimization step (*ρ*_*P*_ = −0.88) compared to other steps (**Fig. 3g,h, Supplementary Figure 7-9**). Intuitively, this strong correlation reflects how likely this test assay can benefit from meta-learning since a smaller loss means the model needs less effort to fit this new assay, allowing end users to identify the assays that can be better improved using ActFound.

### 2.4 A machine learning alternative to FEP calculation

To further illustrate the practical usage of ActFound in drug design, we applied it to two benchmarks for Free Energy Perturbation (FEP) calculation.^37, 42^ FEP calculation is an important computational chemistry approach for real-world drug design by providing the Relative Binding Free Energy (RBFE) between two similar compounds. RBFE reveals how small structure modification impacts the bioactivity of a target protein, guiding the optimization of the drug structure to increase bioactivity.^12^ Nevertheless, FEP calculation is resource-intensive, requiring approximately 24 to 48 GPU hours for computing a single RBFE between two compounds^9, 37^ and is limited to assays with known 3D protein target structure. We hypothesized that RBFE can be approximated by the bioactivity difference in our framework using pairwise learning. To validate this, we collected two benchmarks for FEP from Schrödinger^37^ and Merck.^42^ These benchmarks contain 16 assays and each has 29 compounds on average. Since ChEMBL and BindingDB are collected from diverse sources including scientific articles, there might be data leakage for evaluating FEP benchmarks. To address this, we excluded assays that might lead to data leakage from the training set (see **Methods**), and 58.3% of the compounds on both datasets are unseen in the ChEMBL training set after this processing. We compared the performance of all methods using different proportions of data for fine-tuning ranging from 20% to 80% (**Fig. 4a,b, Supplementary Figure 10-13**). We first observed that our method consistently outperforms competing methods on both datasets using different proportions of fine-tuning data. The improvement over competing methods is larger when fewer data are used, suggesting that our method can more efficiently utilize labeled data in new assays. More importantly, we found that by using 40% of the data for fine-tuning (12 compounds on average for each assay), our model can surpass the performance of FEP calculation tool FEP+(OPLS4),^37^ which is the latest release of the leading commercial software developed by Schrödinger using state-of-the-art OPLS4 force field.^11, 43^ We also compared with the state-of-the-art structure-based methods PBCNet^19^ following their experimental settings, and we found that ActFound outperforms PBCNet even without using any given target protein information (**Supplementary Figure 14**). The promising performance on two FEP benchmarks indicates that our foundation model can serve as a machine-leaning-based alternative to FEP calculations by only requiring a few labeled data points for fine-tuning.

**Figure 4.**
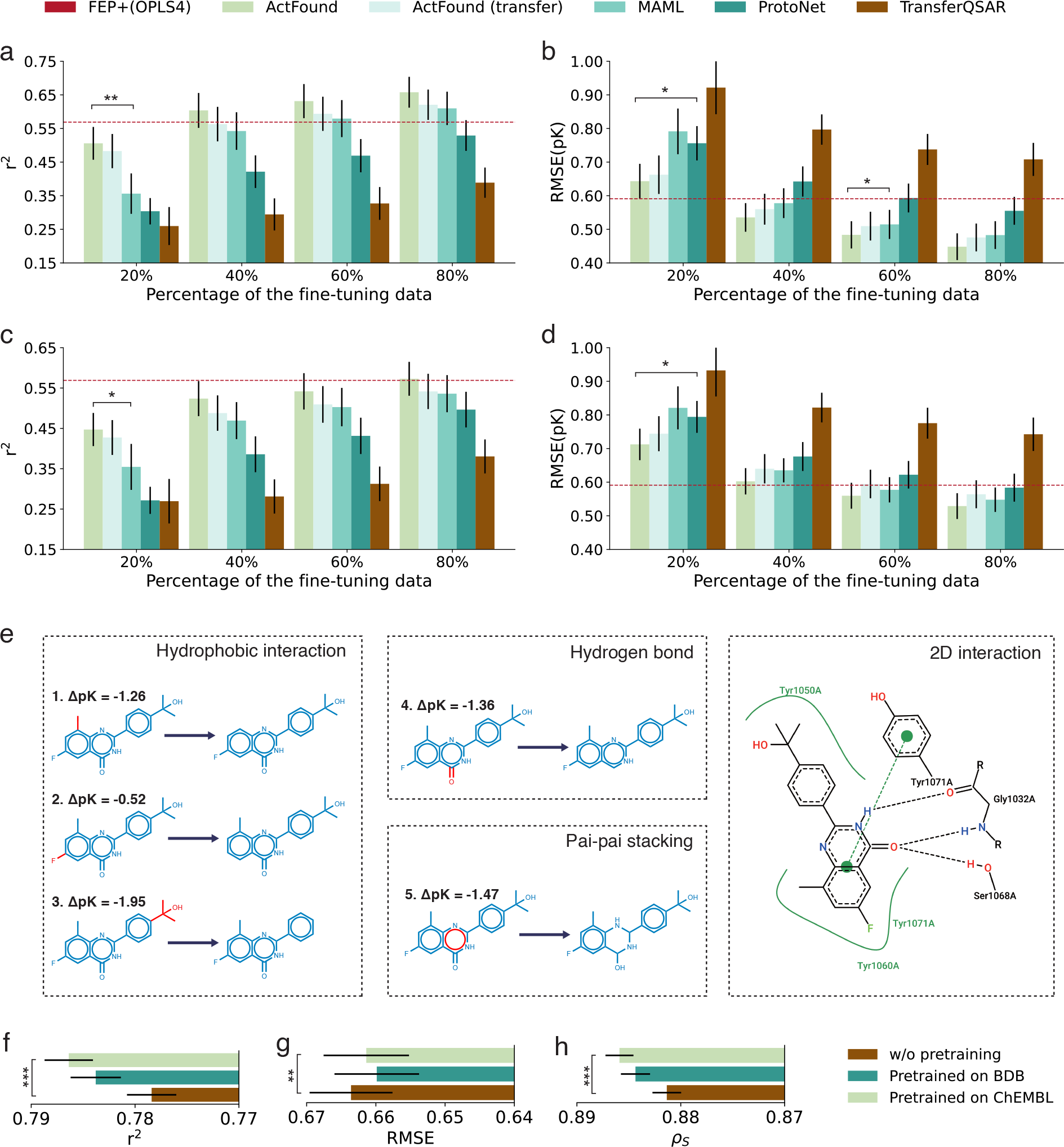
Evaluation of FEP benchmarks and cancer drug response prediction. **a,b**, Bar plots comparing the bioactivity prediction on FEP benchmarks in terms of *r*^2^ (**a**) and RMSE (**b**) when 20%, 40%, 60%, 80% of the experimental bioactivity is used for fine-tuning. All models are trained on ChEMBL. **c,d**, Bar plots comparing the bioactivity prediction on FEP benchmarks in terms of *r*^2^ (**c**) and RMSE (**d**) when the FEP+(OPLS4) calculation results are known for 20%, 40%, 60%, 80% of the fine-tuning compounds. All models are trained on ChEMBL. **e**, Case study on TNKS2, fragments (colored in red) forming non-bonded interactions with the target protein (TNKS2) are removed from the best bioactivity compound. The bioactivity change is predicted using ActFound. **f-h**, Bar plots comparing the performance in terms of *r*^2^ (**f**), RMSE (**g**) and Spearman correlation (**h**) using model pre-trained on BindingDB, model pre-trained on ChEMBL, and randomly initialized model. *ρ*_*s*_ denotes Spearman correlation.

When experimentally measured data are not available, ActFound can alternatively use the data from FEP+(OPLS4) calculation to fine-tune the model. We evaluated this in **Fig. 4c,d**, where we used the results of FEP+(OPLS4) calculation for the fine-tuning compounds and did not incorporate any experimentally measured fine-tuning data. We first observed that the performance of all approaches dropped, indicating that this setting is more challenging than utilizing experimentally measured finetuning data. Nevertheless, ActFound still outperformed all comparison approaches. In contrast to FEP+(OPLS4), we observed that the *r*^2^ of our method is nearly equivalent to that of FEP+(OPLS4) when the fine-tuning compounds contain 80% of the compounds in that assay, and the RMSE value even surpassed FEP+(OPLS4) by 10.4%. Collectively, ActFound can accurately predict bioactivity values in FEP benchmarks when either using experimentally measured data or FEP+(OPLS4) calculated data for fine-tuning, offering a less costly machine learning approach to binding free energy prediction.

Finally, to further understand the prominent performance of ActFound, we presented a case study that illustrates the predicted bioactivity change of different modifications on a given compound. We focused on the compound that has the best bioactivity when TNKS2 is the target protein. In particular, we first identified several potential interactions using Proteins Plus,^44^ then we modified the compound by removing a fragment and predicted the bioactivity value change caused by that modification (**Fig. 4e**). A larger change indicates that the modification is more important. We found that the third compound resulted in the largest bioactivity change (−1.95 pK). This matches the prior chemical knowledge that large non-polar group tends to have strong hydrophobic interactions. We further examined other key interactions by changing the chemical scaffold, which is unseen in the TNKS2 target assay. As the 2D interaction in **Fig. 4e** shows, the best compound forms hydrogen bonds with Gly1032A and Ser1068A, and it also forms pai-pai stacking with Tyr1071A. In the fourth and fifth compounds, the hydrogen bonds and pai-pai stacking are partially eliminated, resulting in a relatively large bioactivity change of −1.36 pK and −1.47 pK respectively, consistent with our expectation. Collectively, this case study reveals how ActFound utilizes pairwise learning to capture the structure difference in functional groups and scaffold, and further enhances bioactivity value prediction.

### 2.5 Cancer drug response prediction

The promising results of ActFound on few-shot bioactivity prediction further motivate us to apply this foundation model to other domains. To this end, we investigated whether ActFound can be used to predict cancer drug response, where the goal is to predict drug sensitivity on new cell lines. Since we can use gene expression values as features for cell lines, it is now possible to test our method in a zero-shot setting where we have never seen any training cell lines for the test drug. We evaluated ActFound on the Genomics of Drug Sensitivity in Cancer (GDSC) dataset.^45^ GDSC contains 969 cell lines, and each cell line has been tested on 207 drugs on average. We concatenated drug features and cell line features to train a supervised model for predicting the sensitivity of the drug to the given input cell line. We compared the performance of using ChEMBL or BindingDB to initialize the drug features against a random initialization (**Fig. 4f-h**). We observed that the ActFound model pre-trained with ChEMBL demonstrated better performance than random initialization on all metrics (p-value < 2×10^−5^), reflecting how ActFound enables the model to learn better compound embeddings. Collectively, the promising result demonstrates the broad applicability of ActFound to drug sensitivity prediction on unseen cancer cell lines.

## 3 Discussion

We have proposed ActFound, a foundation model for compound bioactivity prediction on all assays. ActFound combines meta-learning with pairwise learning to leverage the abundant information in diverse assays and improve the performance on few-shot bioactivity predictions. In particular, we have demonstrated the outstanding performance of ActFound on in-domain bioactivity prediction and cross-domain bioactivity prediction. We also showed the practical usage of ActFound on FEP benchmarks and drug sensitivity prediction for new cell lines. ActFound can serve as the foundation for bioactivity prediction to further promote drug discovery.

ActFound has two limitations that we would like to address in the future. First, although the compound bioactivity prediction approach we developed can be applied to different types of biology assays, it currently cannot consider the meta-data for each assay, such as the sequence of the protein target^46^ or the text assay description^47^ for that assay. We plan to exploit this information to enhance the representation of each assay. Second, ActFound is currently developed using simple molecular fingerprint features without leveraging any pre-trained model for molecular structures. Based on the prominent results of ActFound on all settings we have evaluated, we hypothesize that it can be further improved by incorporating other pre-trained methods. We plan to incorporate pre-trained methods, such as KPGT,^48^ GROVER^49^ and MOLFORMER^50^ to further improve ActFound.

There are a few existing target-free few-shot bioactivity prediction approaches.^20–25^ Compared to these approaches, ActFound has at least two key differences. First, these approaches mainly study bioactivity prediction on assays with molar concentration unit, while ActFound is agnostic to measurement units and is trained on assays of different units. This allows ActFound to generalize to assays with different units. Second, existing approaches predict the absolute value of bioactivity. In contrast, ActFound exploits pairwise learning to predict the relative bioactivity values, which effectively leverages the correlation between bioactivity value rankings of different assays. Pairwise learning has been used for uncertainty quantification on QSAR task,^35, 36^ but has not been applied to few-shot bioactivity prediction. Compared to existing structure-based few-shot binding free energy prediction methods,^17–19^ ActFound also has three key differences. First, these structure-based methods rely on accurate protein-ligand 3D binding pose, which is hard to acquire because of the lack of accurate docking methods. In contrast, ActFound only requires compound information. Second, ActFound exploits meta-learning to enhance the performance on few-shot settings, while existing structure-based methods were mainly based on transfer learning. We showed in our experiments that ActFound outperforms its transfer learning variant. Third, existing structure-based methods were designed specifically for assays with a specific protein target, while ActFound is a general foundation model for compound bioactivity on all assays, which can utilize the large-scale public databases (e.g., ChEMBL, BindingDB) and be applied to assays without any specific protein target, demonstrating the broad applicability of ActFound.

## 4 Method

### 4.1 Problem Setting

We aim to build bioactivity prediction models for different assays in the few-shot setting. Given a limited number of example compounds with known experimental bioactivity for a given assay, ActFound predicts the bioactivity of unseen compounds in that specific assay. We consider bioactivity prediction as a regression task. In our setting, each separate assay serves as an individual task. For the rest of the paper, *p*(*T*) denotes the assay distribution, *T*_*i*_ denotes the *i*-th assay to be trained, 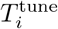 and 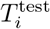 stands for the fine-tuning compounds and test compounds of assay *T*_*i*_, respectively.

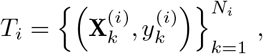

where 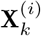 denotes the k-th compound in the assay *T*_*i*_ and 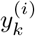 denote its respective bioactivity, *N*_*i*_ stands for the number of compounds in the assay *T*_*i*_. In our experiments, we used 2048-dim Morgan-Fingerprint^51^ extracted by RDKit to represent the compounds **X**.

### 4.2 Review of meta-learning

Meta-learning is a machine learning framework targeting quick adaptation to new tasks. Meta-learning comprises two primary processes: meta-training (also referred to as pre-training) and meta-testing. The model learns a “meta-learner” shared across tasks in the meta-training stage and adapts it to new tasks with a few samples in the meta-testing stage. It is particularly well-suited for bioactivity prediction tasks since there’s a severe shortage of labeled data due to the high costs of laboratory experiments. For instance, only 12.7 compounds are measured for each assay in ChEMBL^7^ on average.

We adopt MAML, a widely-used meta-learning framework, for the training of our bioactivity prediction model ActFound. MAML aims to obtain optimal initial parameters *θ*^*^ shared across all assays. It achieves this through an iterative bi-level optimization process during the meta-training phase, consisting of an inner loop and an outer loop. In the inner loop, the model is rapidly adapted to the fine-tuning compounds 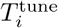 over a small number of iterations, to make the model’s parameters suitable for the particular assay *T*_*i*_. In the outer loop, the model’s parameters are updated based on the performance of the adapted model in the test compounds of the assay 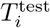, which aims to make the model’s initial parameters more suitable for fast adaptation across various assays.

Specifically, in the inner loop stage, MAML samples a set of assays *T*_*i*_ ∼ *p*(*T*) and adapts model parameters *θ* to obtain assay-specific parameters 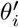 using the following equation:

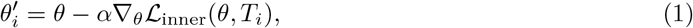

where *α* is the inner loop learning rate hyper-parameter, and ℒ_inner_(*θ, T*_*i*_) is the the inner loop loss. In the outer loop stage, MAML obtains the outer loop loss 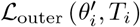 in the test compounds 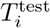 for each training assay *T*_*i*_ ∼ *p*(*T*). It then updates the model parameters *θ* using the outer loop loss based on the following equation:

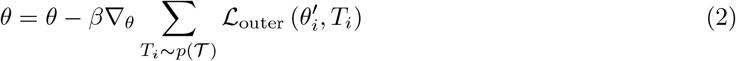

where *β* is the outer loop learning rate hyper-parameter.

After the meta-training stage, we get optimal initial parameters *θ*^*^ suitable for fast adaptation on various assays. In the meta-testing stage, MAML only takes the inner loop stage for an unseen test assay *T*_*j*_ following Equation 1. Subsequently, MAML provides predictions for the test assay *T*_*j*_ using the model parameters 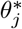, which is fine-tuned from *θ*^*^ on the test assay’s fine-tuning compounds 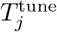.

### 4.3 Pairwise learning

Since there are many types of assays with varying units (e.g., nMol, mg/kg/day, ug/ml), value ranges, and measurements, directly learning the absolute bioactivity among assays may encounter issues of incompatibility among assays, which can subsequently result in poor transferability to unseen assay types and harm overall performance. ActFound aims to predict the relative bioactivity difference between compounds’ pairs and then construct the predicted bioactivity for the test compounds of an assay, which makes the model agnostic to assay units and value ranges. It adopts the Siamese Network architecture,^18^ where the input is a pair of compounds, and the prediction is the bioactivity difference between them.

Specifically, for the k-th compounds 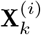 and the j-th compounds 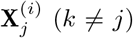 in the *i*-th assay *T*_*i*_ ∼ *p*(*T*), the predicted bioactivity difference 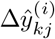 between the two compounds is computed as:

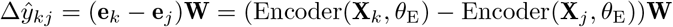

where **e**_*k*_, **e**_*j*_ are the embeddings of the two compounds. *θ*_E_ is the parameters for the compound encoder, which is implemented as a 2-layer MLP with the hidden size of 2, 048. **W** ∈ ℝ^2048×1^ is the weight matrix for the last linear layer, and the model parameters of ActFound can be defined as *θ* = (*θ*_E_, **W**). For reduced computation consideration, we followed ANIL^52^ to take the fast adaptation method, which only optimizes the last linear layer parameters **W** in the inner loop. So the assay-specific parameter 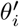 adapted for assay *T*_*i*_ after inner loop should be equal to 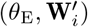.

As the Siamese Network model only outputs the relative bioactivity difference between two compounds, we further need to construct the absolute bioactivity of the test compounds 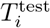 in the assay *T*_*i*_. For each fine-tuning compound 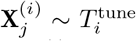, we can get a constructed bioactivity for 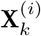 using the equation 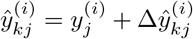, where 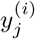 is the experimental bioactivity of 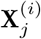. So there is at most 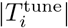 constructed absolute bioactivity for 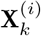. After this, we utilized the attention mechanism^53^ with the Tanimoto mask to fusion them together. Our intuition is that similar compounds usually have similar bioactivities (this is why KNN works for bioactivity prediction^54^), thus lowering the absolute error of the constructed bioactivity. So similar compounds should contribute more to the final prediction 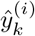. The equation for the final prediction 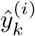 of 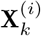 is given as follows:

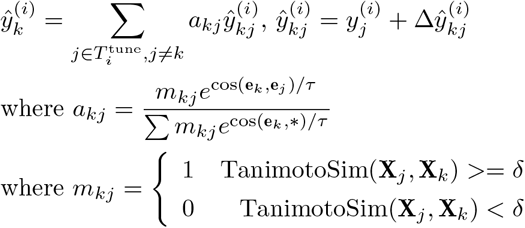

where cos(**e**_*k*_, **e**_*j*_) is the cosine similarity between embeddings of the compound 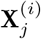 and 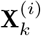, *τ* is the temperature hyper-parameter for attention, and *δ* is a threshold hyperparameter, which is implemented as the median of Tanimoto Similarity values between 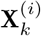 and all other fine-tuning compounds 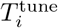. The Tanimoto mask *m*_*kj*_ effectively filters out the influence of dissimilar compound pairs, which typically exhibit large absolute errors. This filtering mechanism has been demonstrated to enhance the overall performance of ActFound.

The training objective for ActFound, both in the inner loop and the outer loop, encompasses two key components: pairwise loss and absolute loss. For the pairwise loss, it is characterized as the mean square loss, which is calculated between the predicted relative bioactivity 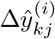 and the experimental relative bioactivity 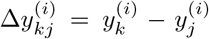. On the other hand, the absolute loss is simply the mean square loss between the constructed bioactivity 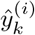 and the experimental bioactivity 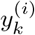. It’s worth noting that the difference between the inner loop loss ℒ_inner_ and the outer loop loss ℒ_outer_ is minimal. For the inner loop loss, the pairwise loss is defined on the compound pairs within the fine-tuning compounds 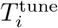, and the absolute loss is similarly defined on 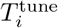. For the outer loop loss, the pairwise loss is defined on the compound pairs formed between the fine-tuning compounds 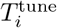 and the test compounds 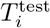 along with that formed within the test compounds, while the absolute loss is solely defined on 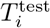.

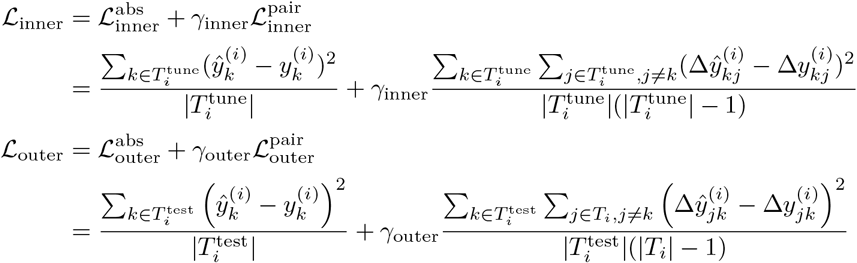

where *γ*_inner_, *γ*_outer_ are two hyper-parameters for the pairwise loss weight. Note that ℒ_inner_ can be written in ℒ_inner_ (*θ, T*_*i*_) = ℒ_inner_ (*θ*_E_, **W**, *T*_*i*_), and ℒ_outer_ can be written in 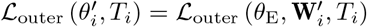.

### 4.4 KNN-MAML and fusion method

Many bioactivity assays exhibit similarities, such as those assessing the binding affinity to both the wild type and mutant forms of a specific target protein. This observation inspired us to leverage the information from similar assays in the training set to improve testing on previously unseen assays. We proposed a novel algorithm called *k*-Nearest Neighbors MAML (KNN-MAML), which dynamically identifies similar assays during the meta-testing process and utilizes them to aid the adaptation of unseen assays.

In the inner loop of ActFound for each test assay, we searched for *k*-similar assays from the training set, and adapted the trained initial parameters *θ*^*^ to the assay-specific parameters 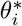, using the data from these *k* assays.

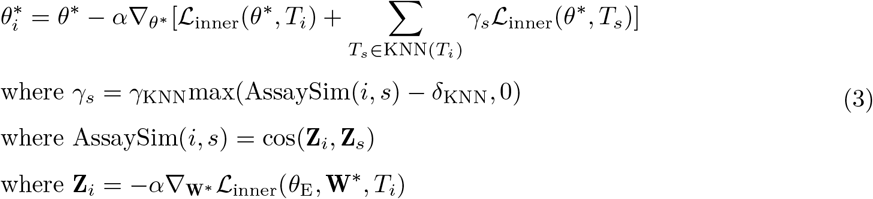

where *γ*_*s*_ is the weight of the assay *T*_*s*_ in *k*-similar assays KNN(*T*_*i*_), AssaySim(*i, s*) is the assay similarity between *T*_*i*_ and *T*_*s*_, **Z**_*i*_ is the assay feature for *T*_*i*_, and *γ*_KNN_, *δ*_KNN_ are two hyper-parameters controlling the contribution of each *k*-similar assays. As stated above, we employed the fast adaptation method,^52^ which only optimizes the last linear layer parameters **W**^*^ in the inner loop. As we assume that similar assays should have similar assay-specific model parameters, we can straightforwardly define the assay feature **Z**_*i*_ as the gradient of the parameters **W**^*^ with respect to the inner loop loss, which is also identical to 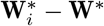. With this definition in place, we measure the similarity between assays as the cosine similarity between two assay features cos(**Z**_*i*_, **Z**_*s*_).

In addition to the KNN-MAML method, we also developed a straightforward ensemble approach for combining ActFound and its transfer learning variant ActFound(transfer). We observed a significant correlation between the inner loop loss for the first optimization step ℒ_inner_(*θ, T*_*i*_) (we performed 5 inner loop optimization steps) and the test performance of the test assay (**Fig. 3g,h, Supplementary Figure. 7-9**). Building on this insight, we propose to fusion ActFound and ActFound(transfer) based on the loss of the first inner loop step. Specifically, we calculated the first-step loss for both ActFound and ActFound(transfer) on the test assay. When the loss difference between ActFound and ActFound(transfer) exceeds a threshold (typically −0.1), we combine the results of the two methods; otherwise, we solely employ ActFound. This straightforward approach has demonstrated improved performance across most tests, providing end users with a robust strategy for determining when to utilize ActFound.

### 4.5 Implementation details

The training strategy of ActFound mainly follows MAML++,^55^ and the details are listed as follows. In the meta-training (pre-training) stage, ActFound are trained for 60 epochs on ChEMBL and BindingDB, and 80 epochs for FS-MOL and pQSAR-ChEMBL. For the inner loop, we utilized a simple stochastic gradient descent (SGD) optimizer, we performed 5 steps with separate batch normalization parameters for each step. The learning rates for each step of the inner loop are also learnable parameters, which means that the model will update the learning rate parameters during the meta-training stage. For the outer loop, we used the Adam optimizer with a cosine annealing learning rate scheduler, with the maximum learning rate to be 0.00015 and the minimum to be 0.0001. The remaining parameters for the Adam optimizer were configured with the default settings of PyTorch. We applied early stopping for all methods and utilized the best validation epoch for testing. We set the meta-batch size to 16 for all methods, which means each meta-batch contains 16 assays.

### 4.6 Comparative methods and evaluation metrics

In our experiment, we implemented several transfer learning and meta-learning comparative methods, including ActFound(transfer) (transfer learning variant of ActFound), MAML,^38^ DKT (Deep kernel transfer),^56^ CNP (Conditional neural processes),^57^ ProtoNet^39^ and TransferQSAR (transfer learning variant of MAML), and the transfer learning variant means that we didn’t take the inner loop during training. When testing on new assays, we fine-tuned the last linear layer of transfer learning methods (ActFound(transfer) and TransferQSAR) on the test assay, which is implemented as a 5 steps optimization using the SGD optimizer. The fine-tuning learning rate was 0.004, which is the optimal setting determined in our grid search on the validation assays. For fairness consideration, We applied an early stopping strategy for all methods on the validation assays, and they have the same hyper-parameter settings. All methods share the backbone model (3-layer MLP) and the same inputs (2048-dim Morgan fingerprint) which is identical to ActFound. The hyper-parameters (including learning rate and its scheduler, training epoch, meta-batch-size, and inner loop and outer loop optimizer parameter) are also identical to ActFound. Except for the ADKF-IFT^23^ model on the in-domain experiment of FS-MOl, we directly used the released checkpoints for testing. The ADKF-IFT model is a combination of fingerprint-based and GNN-based models, which takes the input of 2048-dim Morgan fingerprint and 2D molecular graph. Three single-task methods, including RF (Random Forest),^58^ KNN (*k*-Nearest Neighbors)^59^ and GPST (single-task GP with Tanimoto kernel)^60^ also take 2048-dim Morgan fingerprint as an input.

We employed two metrics for evaluation, *r*^2^ and RMSE. *r*^2^ is defined as *max*(*ρ*_*P*_, 0)^2^, where *ρ*_*P*_ is the Pearson correlation between the predicted and experimental bioactivities. RMSE means the root mean square error between the predicted and experimental bioactivities on the test compounds of the test assay. Unless otherwise specified, all test experiments were repeated for 10 times, and the fine-tuning compounds of the test assay were uniformly split at random.

### 4.7 Training data curation

Our data were curated from two publicly available datasets: ChEMBL^7^ and BindingDB.^14^ For ChEMBL dataset curation, we downloaded the ChEMBL32 dataset from the official website. The original ChEMBL32 dataset contains 2.4 million unique molecules and 1.6 million manually curated assays which come from scientific literature, patents, and other sources. All assays in ChEMBL32 are meticulously categorized, encompassing various aspects of the real-world drug design. After careful curation, our processed ChEMBL dataset contains 35, 644 assays, 1.4 million tested bioactivities, and 0.7 million unique compounds, with each assay containing 39.3 compounds on average. All assays have either a molar concentration unit (e.g. nMol), density unit (e.g. ug/mL), or dosing unit (e.g. mg/kg/day). We hypothesize that this choice could maximize the assay diversity while ensuring the quality.

The curation pipeline for ChEMBL mainly contains three stages. First, all candidate assays with the required number of tested bioactivities (≥ 20) and desirable BAO (Bio-Assay-Ontology) format are collected from ChEMBL32, and we kept all bioactivities with molar, density, or dosing unit. As we aim to predict bioactivity for drug-like small molecules, in the second step, we cleaned all compounds in our curated data by removing all inorganic and large molecules with molecular weights larger than 1, 000. Third, further cleaning of test bioactivity data for all assays was implemented. Specifically, we removed duplicate tested bioactivity measurements of one compound by simply taking the results’ average. We removed all bioactivity with the standard relation not equal to “=”, which means that its accurate value is unknown and might contain some noise. We also removed all “orphan compounds” that are dissimilar to all other compounds in this assay, which means that all other compounds have Tanimoto similarity smaller than 0.2 with it.

For BindingDB dataset curation, the original BindingDB version 2022m7 was downloaded from https://www.bindingdb.org, which contains 1.2 million compounds and 9.2 thousand targets. Our curated BindingDB dataset contains 15, 225 assays, 0.9 million tested bioactivities, and 0.4 million unique compounds, and each assay contains 60.9 compounds on average. The curation pipeline for BindingDB is almost identical to ChEMBL’s curation. However, since all assays in BindingDB had IC50, Ki, Kd, or EC50 measurements which already met our requirement, we didn’t implement the first curation stage for BindingDB. The BindingDB dataset only contains assays with molar concentration units, which means that the model trained on BindingDB might have a relatively poor transfer ability to assays of other units.

### 4.8 In-domain experiment setting

For all in-domain experimental settings, we first trained models on the training assays for each dataset and then tested them on the test assays of the corresponding dataset. All in-domain test experiments were repeated for 10 times, and the fine-tuning compounds of each test assay were uniformly split at random. However, the in-domain experimental on pQSAR-ChEMBL is only conducted once because assays on pQSAR-ChEMBL follow a fixed scaffold split. We implemented in-domain experiments for four datasets: our curated ChEMBL and BindingDB datasets (see Section 4.7 for more details), together with two other public datasets: FS-Mol^22^ and pQSAR-ChEMBL,^21^ which is curated from the earlier version of ChEMBL. The FS-Mol dataset contains 4, 938 assays, 0.4 million tested bioactivities, and 0.2 million unique compounds, each assay contains 86.3 compounds on average. The pQSAR-ChEMBL dataset contains 4, 276 assays and 0.5 million unique compounds, and the average number of compounds in each assay is 320.0.

For the in-domain experiment on ChEMBL and BindingDB datasets, we split out a total of 1000 assays uniformly at random to prepare our validation and test assays (500 assays as validation and 500 assays as validation respectively). We performed few-shot (16-shot) testing on the test assays of these two datasets, which means that the model is fine-tuned on the 16 fine-tuning compounds of the test assay, and subsequently tested on the remaining test compounds.

For the in-domain experiment on FS-Mol, assay split is referenced from the setting of Chen et al,^23^ which has 40 assays for validation and 111 assays for test respectively. We performed 16, 32, 64, 128 shot testing for all 111 test assays on FS-Mol.

For the in-domain experiment on pQSAR-ChEMBL, we adopted the assay split configuration of Yao et al.^61^ This setup consists of 76 assays for validation and 100 assays for testing. The assays on pQSAR-ChEMBL follow the scaffold split setting,^21^ where the fine-tuning and the test compounds of an assay don’t have any overlapping chemical scaffold. For each assay in pQSAR-ChEMBL, roughly 75% of the compounds are used for fine-tuning, with the remaining compounds assigned to the test compounds.

For Activity-ChEMBL testing, we manually curated 178 assays with the assay type equal to “Activity” and percentage unit (%). Each assay has 32.6 compounds on average. The Activity-ChEMBL set can also be viewed as an out-of-domain set, as all bioactivities have percentage units, which is not included in our curated ChEMBL dataset. However, unlike other datasets, we don’t know whether a higher raw value corresponds to better or worse bioactivity for assays in the Activity-ChEMBL dataset. To solve this problem, we ran ActFound and ActFound(transfer) two times during testing, which means that the model is fine-tuned twice to predict the positive and the negative relative bioactivity respectively. After that, we took a simple fusion approach to select one of those two independent running results, and we employed the first step loss for selection, which again utilized our observation of the high correlation between loss for the first fine-tuning step and the test performance (**Fig. 3g,h, Supplementary Figure. 7-9**).

### 4.9 Cross-domain experiment setting

In cross-domain experiments, we initially trained models on the ChEMBL and BindingDB datasets and subsequently applied them for testing on other datasets. The cross-domain test experiments were repeated 10 times. Similar to in-domain experiments, the fine-tuning compounds of a test assay were also uniformly split at random.

We first performed ChEMBL-to-BindingDB and BindingDB-to-ChEMBL experiments. However, as ChEMBL and BindingDB are both collected from sharing sources (including public journals, patents, and other databases), they may have lots of overlapping assays. Given that both the test assays of ChEMBL and BindingDB are uniformly split at random, there is a risk that assays in the BindingDB test set may have already appeared in the ChEMBL training set. To mitigate the risk of data leakage caused by these overlapping assays in the ChEMBL-to-BindingDB and BindingDB-to-ChEMBL cross-domain test, we adopted the approach outlined in pQSAR-ChEMBL^21^ to find identical assays and then remove them from the test assays. Specifically, if the bioactivity Pearson correlation (*ρ*_*P*_) between the shared compounds of two assays exceeded 0.99, we considered those two assays as identical. For each assay in the BindingDB test set, if we found an identical assay in ChEMBL, we removed the specific test assay from the test set. Following this data leakage prevention procedure, we retained 203 assays for the ChEMBL-to-BindingDB test and 351 assays for the BindingDB-to-ChEMBL test.

We also studied the transfer ability of the models on two kinase inhibitor datasets: KIBA^40^ and Davis,^41^ here we performed 16-shot testing for Davis, and 16, 32, 64, 128-shot testing for KIBA. Both datasets are directly downloaded from https://github.com/thinng/GraphDTA.^62^ The Davis dataset contains 30, 056 Kd measured binding affinities (one kind of bioactivity) of 68 compounds and 442 kinases, and the KIBA dataset contains 118, 254 KIBA scores of 2, 111 compounds and 229 kinases (each kinase can be roughly considered as an assay). The KIBA dataset was initially created by merging diverse assays with IC50, Ki, and Kd measurements, ultimately consolidating them into a unified KIBA score, and higher KIBA scores indicate better bioactivity. Notably, the KIBA score is not present in the ChEMBL and BindingDB datasets, which is the main difference between KIBA and Davis. Similar to the in-domain experiments, we only retained assays with more than 20 compounds for testing in the KIBA and Davis datasets. After this data preprocessing, the final KIBA dataset contains 165 kinases and 2,068 unique compounds, with an average of 617.3 compounds per target. The Davis dataset contained 212 kinases, 68 unique compounds, and an average of 29.6 compounds per target.

As Davis may share common data sources with ChEMBL and BindingDB, there also might be a data leakage problem in Davis. To address this potential data leakage issue in the Davis dataset, we carefully removed any assays that could lead to data leakage from the ChEMBL and BindingDB training sets. To achieve this, we detected all assays in ChEMBL and BindingDB which have at least one identical assay in Davis, and then we removed them from the training set. We again followed pQSAR-ChEMBL^21^ for detecting identical assays.

### 4.10 FEP benchmark experimental setting

We evaluated the performance of trained models on two well-known FEP calculation benchmarks from Schrödinger^37^ and Merck.^42^ The FEP benchmark contains 16 assays, with an average of 28.8 compounds for each assay. Given the relatively small number of assays in the FEP set, we conducted test experiments for 40 times, and the fine-tuning compounds for test assays were randomly and uniformly split.

Since the ChEMBL and BindingDB datasets both contain data sources from public journals, there’s a possibility that assays in the FEP set overlap with those in ChEMBL or BindingDB. To address this data leakage issue, we systematically identified all assays in ChEMBL and BindingDB that had at least one identical assay in the FEP set. Subsequently, we manually removed these overlapping assays from the training sets. Similar to the approach used in cross-domain experiments, the method for detecting identical assays was adopted from pQSAR-ChEMBL.^21^ After data leakage processing, 58.3% of the compounds in the FEP set are unseen in ChEMBL. Data for the FEP set assays and the FEP+(OPLS4)^37^ calculated results were downloaded from https://github.com/schrodinger/public_binding_free_energy_benchmark,^43^ we directly read the compounds’ smiles from the SDF files, and manually deleted all formal charges.

For our case study on the TNKS2 target assay, we first fine-tuned ActFound on nearly all compounds in the assay and then predicted the bioactivity of modified compounds. Note that we removed all modified compounds from fine-tuning compounds, to avoid the potential data leakage problem in this study. The protein structure for detecting the interaction graph is taken from Protein Data Bank(PDB) using PDB id 4UI5, and the interaction graph is created using Proteins Plus.^44^

### 4.11 Cancer drug response experimental setting

For experiments on the GDSC dataset, we downloaded the Genomics of Drug Sensitivity in Cancer (GDSC)^45^ dataset from the official website. The dataset contains 621 drugs, 969 cell lines, and 243, 464 drug-cell-IC50 triplets. We obtained the SMILES of drugs using PubChemPy and Rdkit packages and ignored all drugs that failed to get valid SMILES. Following these preprocessing steps, the refined GDSC dataset included 969 cell lines, with an average of 206.5 drugs per cell line. We employed a 425-dimensional mutation feature as the representation for cell lines. The cell line branch model is a simple 2-layer MLP with the hidden size set to 64 and the output dimension set to 2,048. The output vector from the cell line branch is further processed through a sigmoid layer, followed by fusion with the compound embedding e_*k*_ using a dot product operation. This fused representation is ultimately passed through a linear layer to predict drug sensitivity. In the GDSC experiment, we split 100 cell lines for training from the GDSC dataset, and the other 869 cell lines for testing. We simply trained all models for 80 epochs and then reported the best test result among all epochs. During the testing stage, we took the zero-shot setting, which means that we simply predicted the drug sensitivity of compounds to the input cell line without any fine-tuning.

### 4.12 Bioactivity definition

The definition of bioactivity varies depending on the type of assay being used. For all assays with molar concentration unit (e.g. nMol), density unit (e.g. ug/mL), dosing unit (e.g. mg/kg/day), and percentage unit (%), the bioactivity is defined as the negative log of the raw value pX = − log_10_ X, where X is the raw value in the original dataset. The bioactivity in most of the dataset we used are defined in this way (including ChEMBL, ChEMBL-Activity, BindingDB, FS-MOL, pQSAR-ChEMBL, Davis, GDSC). For assays on KIBA, we directly employed the KIBA score as the bioactivity. For assays on the FEP set, the direct downloaded data are binding free energy with kcal/mol unit, and the lower binding free energy indicates better bioactivity. For consistency with other assays in this paper, we converted them to the negative log of the dissociation constant pK = −log_10_K using Gibbs free energy isotherm equation, where K means the dissociation constant. The conversion equation can be simply written as ΔG = −1.363pK under the temperature of 298.15k.

## Code and data availability

The data and checkpoints of ActFound and other comparative methods (ActFound(transfer), TransferQSAR, ProtoNet^39^ and MAML^38^) are available in the Google Drive https://figshare.com/articles/dataset/ActFound_data/24452680. The code is available in our GitHub project https://github.com/BFeng14/ActFound.

## Acknowledgments

We’d like to express our gratitude to Dr. Michael K. Gilson and Dr. Tiqing Liu from the University of California, San Diego (UCSD) for their advice on data curation. Additionally, we’d like to express our gratitude to Dr. Fenglei Cao from Inf Technology for his guidance on the FEP benchmark experiments.

## Supplementary Information

**Supplementary Figure 1.**
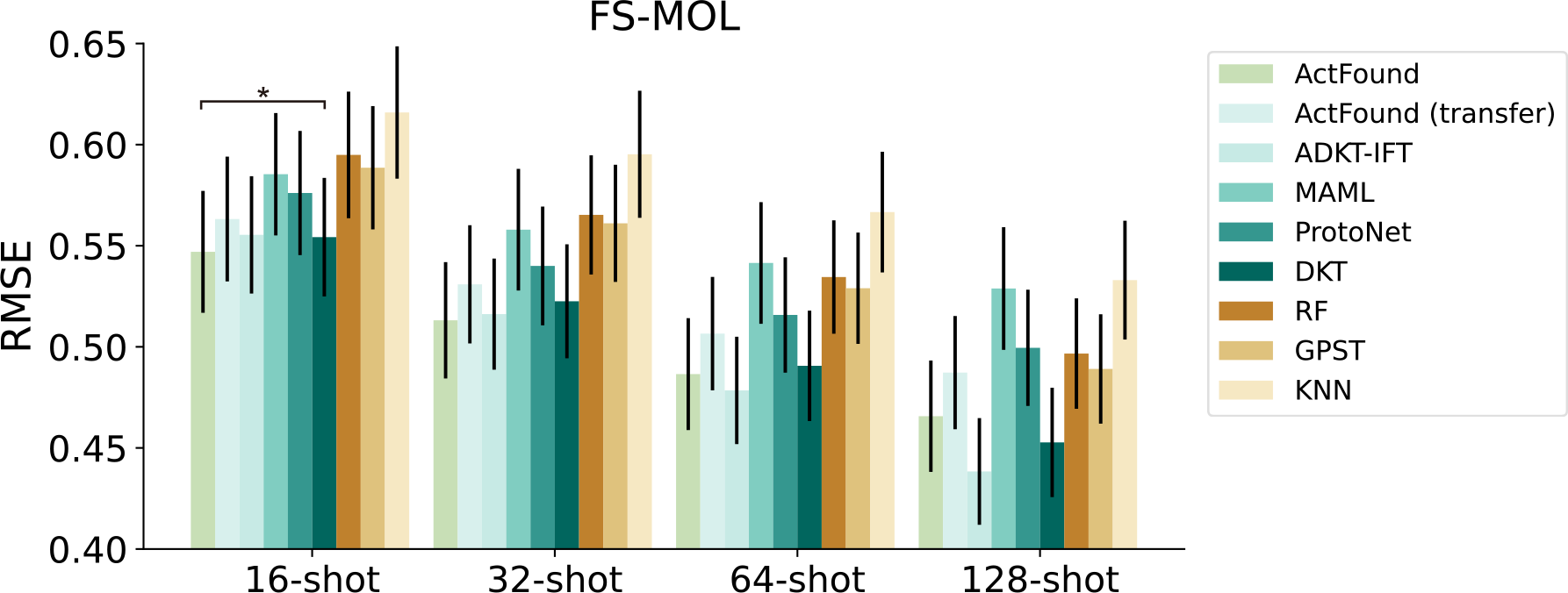
Bar plot comparing the 16, 32, 64, 128-shot bioactivity prediction on FS-MOL in terms of RMSE. The asterisks indicate that ActFound outperforms the next best-performing model in the metric, with t-test significance levels of p-value < 5 × 10^−2^ for *, p-value < 5 × 10^−3^ for **, and p-value < 5 × 10^−4^ for ***.

**Supplementary Figure 2.**
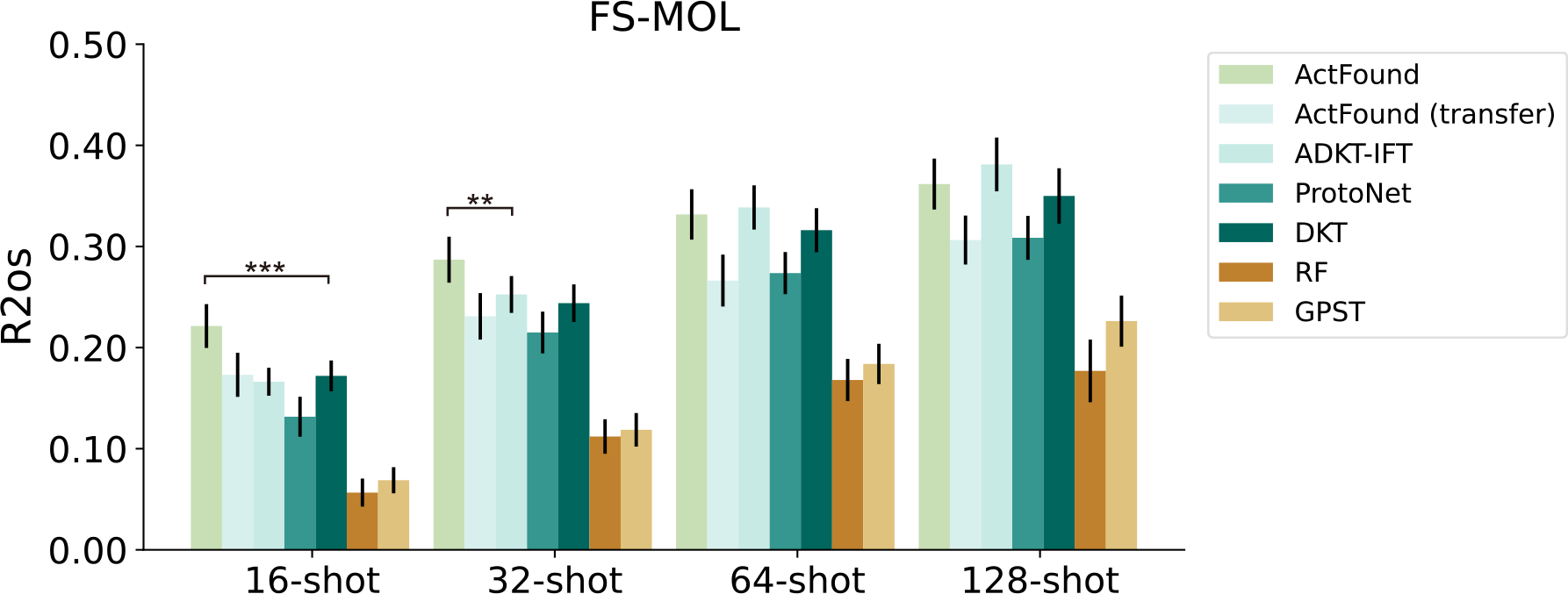
Bar plot comparing the 16, 32, 64, 128-shot bioactivity prediction on FS-MOL in terms of R2os. We adopted the definition of R2os from ADKF-IFT.^23^

**Supplementary Figure 3.**
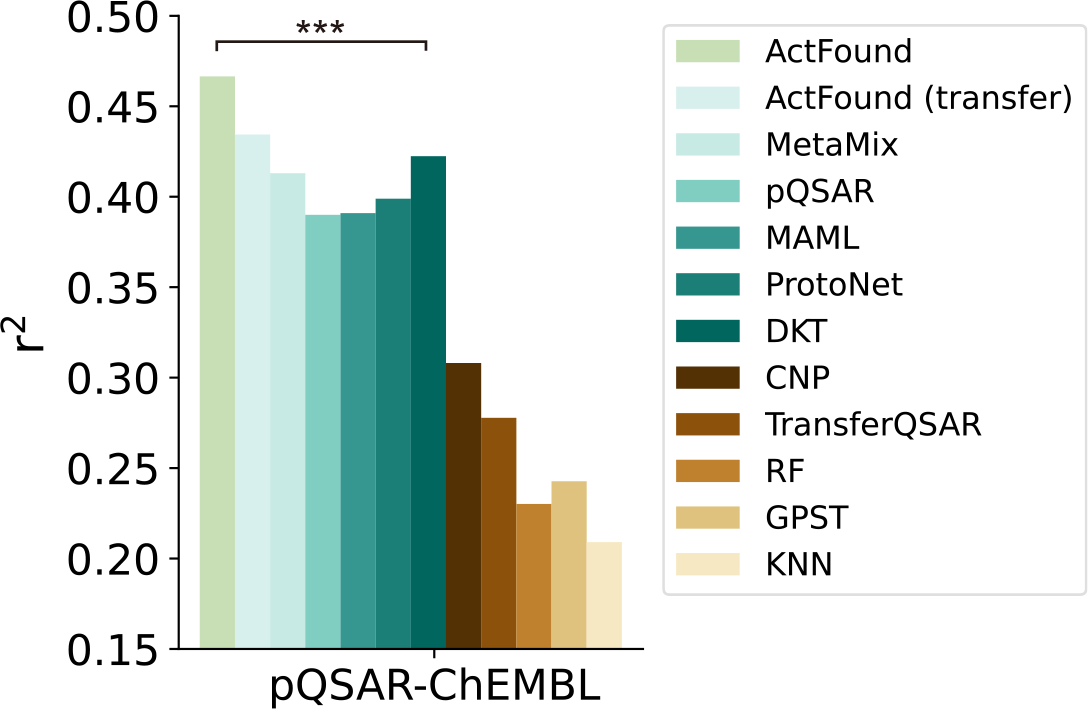
Bar plot comparing bioactivity prediction using scaffold data splitting on pQSAR-ChEMBL in terms of *r*^2^. 75% of the experimental bioactivity is used for fine-tuning. The results of MetaMix and pQSAR are collected from the paper of MetaMix.^61^ MetaMix is a meta-learning method. pQSAR is a multi-task learning method developed by Novartis.^21^

**Supplementary Figure 4.**
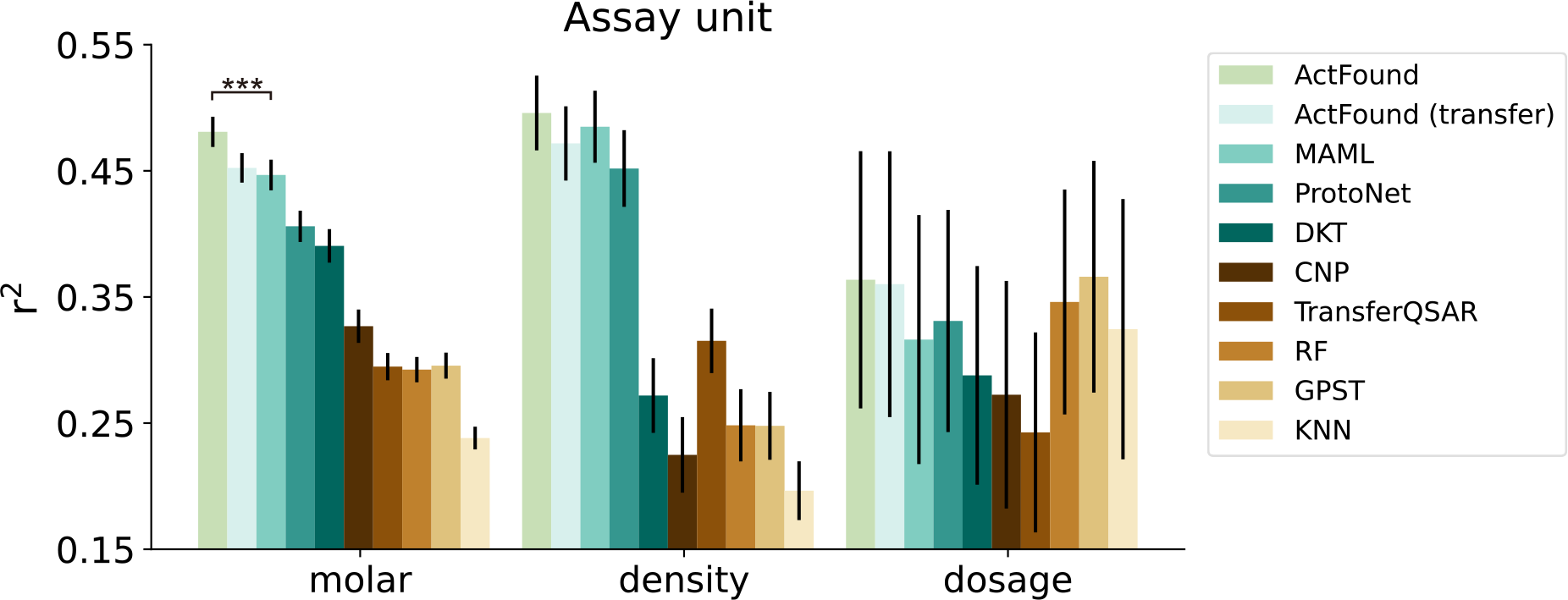
Bar plot comparing ActFound and competing methods in 16-shot bioactivity prediction on ChEMBL in terms of *r*^2^. Assays of molar concentration unit (e.g. nMol), density unit (e.g. ug/mL), and dosing unit (e.g. mg/kg/day) in the test assays of ChEMBL are evaluated separately.

**Supplementary Figure 5.**
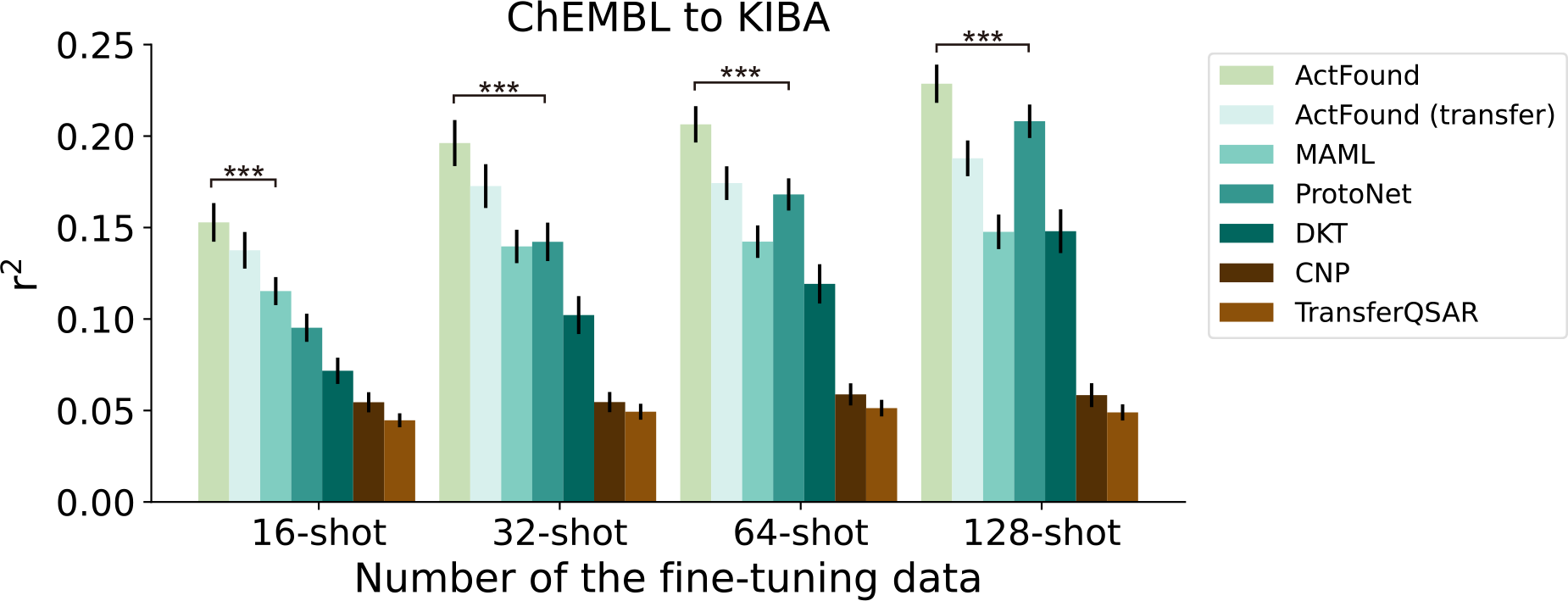
Bar plots comparing the 16, 32, 64, 128-shot bioactivity prediction of ChEMBL-to-KIBA in terms of *r*^2^.

**Supplementary Figure 6.**
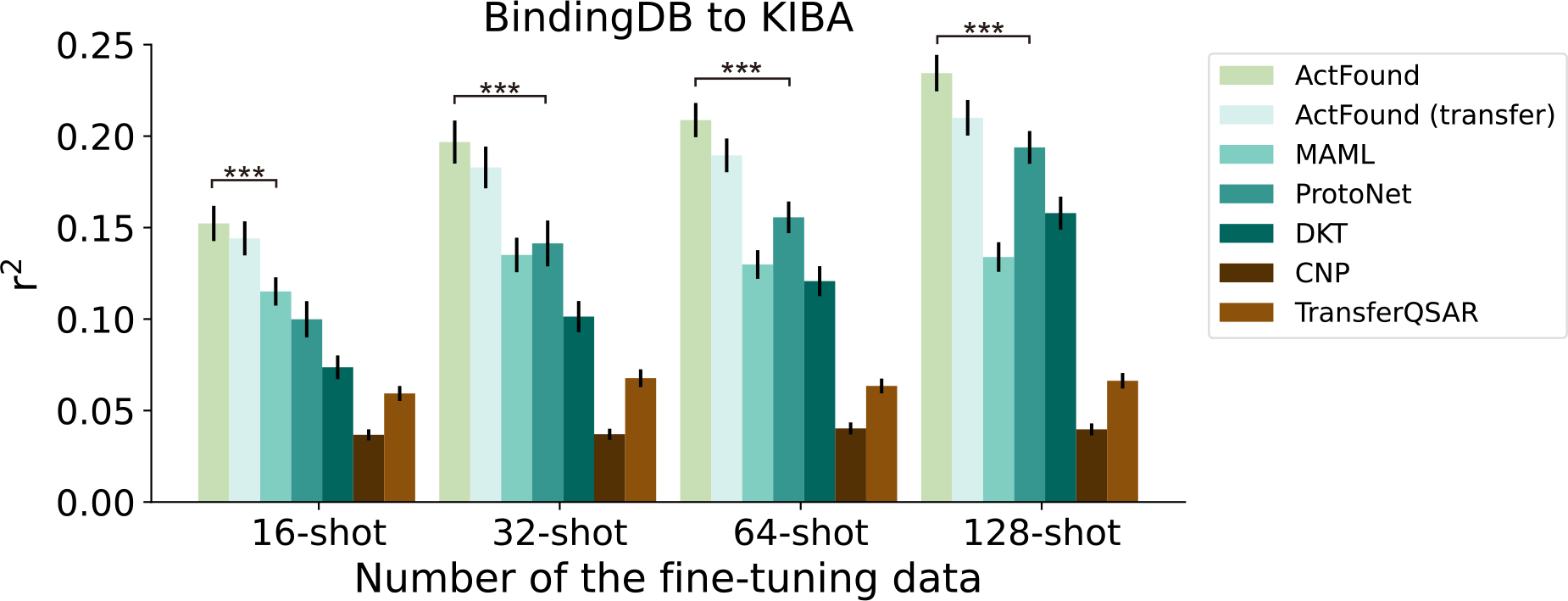
Bar plots comparing the 16, 32, 64, 128-shot bioactivity prediction of BindingDB-to-KIBA in terms of *r*^2^.

**Supplementary Figure 7.**
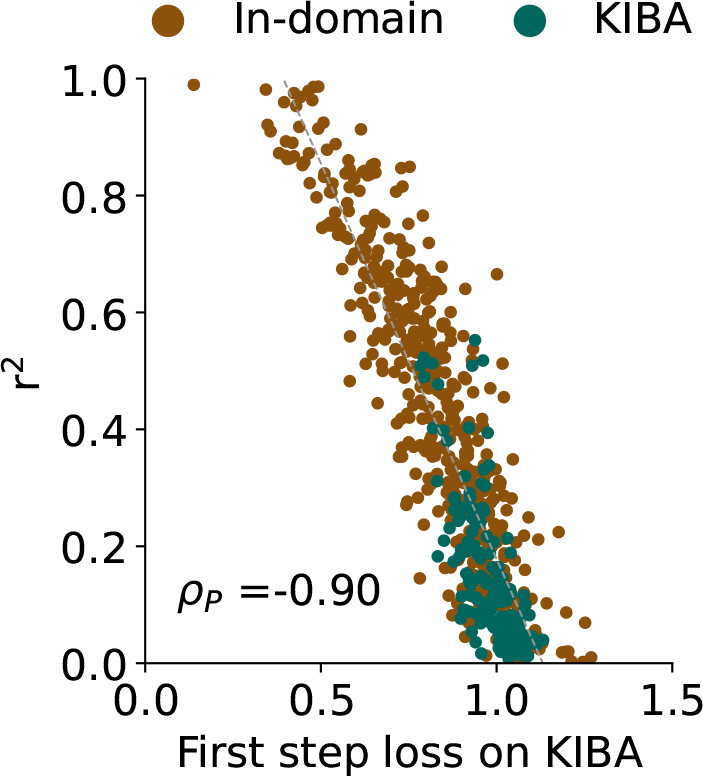
Scatter plots comparing the loss for the first step and the test performance in terms of *r*^2^ on the BindingDB-to-KIBA cross-domain experiment. The test performance (*r*^2^) of the test assay is strongly correlated with the loss for the first step in terms of Pearson Correlation.

**Supplementary Figure 8.**
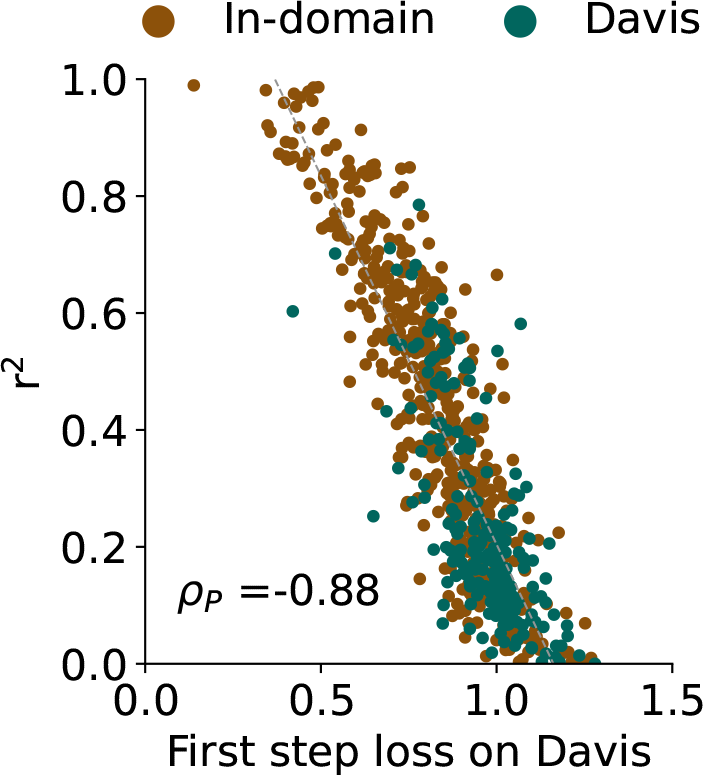
Scatter plots comparing the loss for the first step and the test performance in terms of *r*^2^ on BindingDB-to-Davis cross-domain experiment. The test performance (*r*^2^) of the test assay is strongly correlated with the loss for the first step in terms of Pearson Correlation.

**Supplementary Figure 9.**
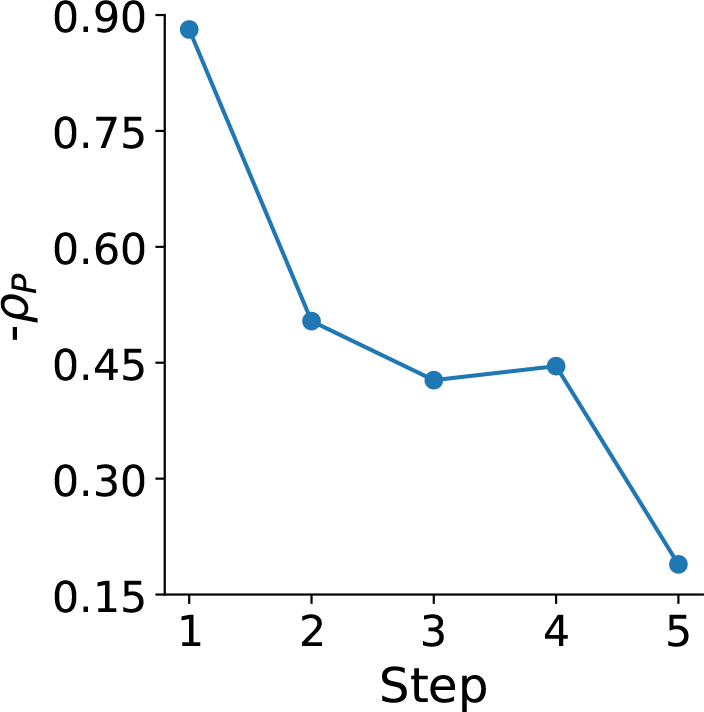
Negative Pearson correlations (*y*-axis) between the loss after the *k*-th step (*x*-axis) and the test performance (*r*^2^) after convergence.

**Supplementary Figure 10.**
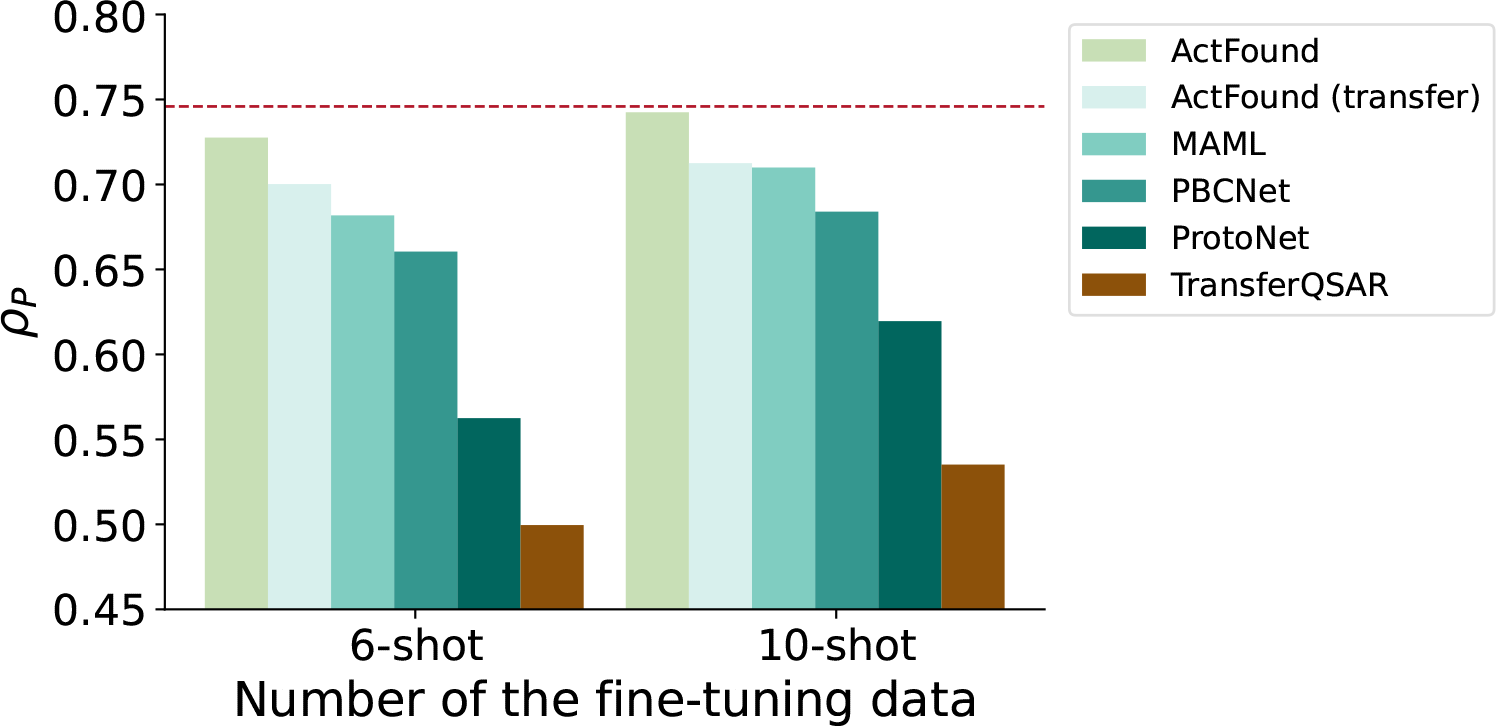
Bar plot comparing the 6-shot and 10-shot bioactivity prediction on FEP benchmarks in terms of *ρ*_*P*_. The result of PBCNet is collected from its original paper.^19^ The red dashed line represents the computational results of FEP+(OPLS4). The Thrombin assay is ignored in the 10-shot test as it only contains 11 compounds.

**Supplementary Figure 11.**
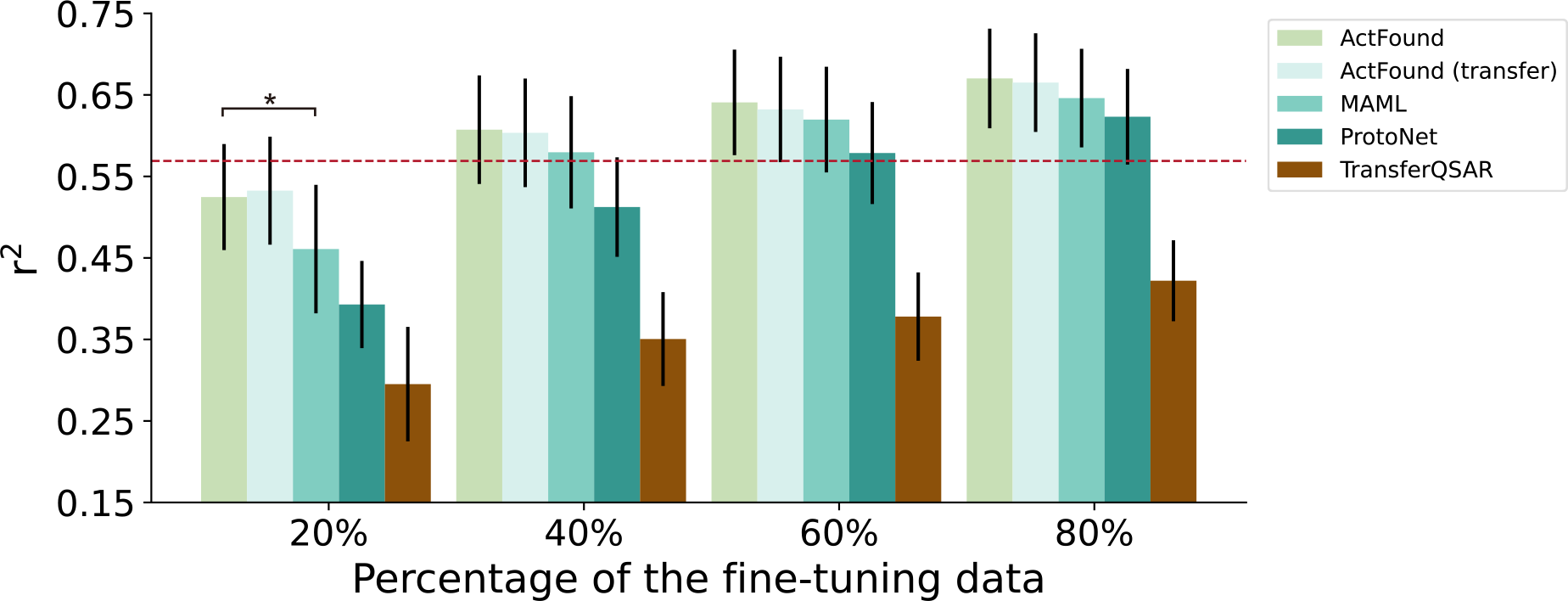
Bar plot comparing the bioactivity prediction on FEP benchmarks in terms of *r*^2^ when 20%, 40%, 60%, 80% of the experimental bioactivity is used for fine-tuning. All models are trained on BindingDB. The red dashed line represents the computational results of FEP+(OPLS4).

**Supplementary Figure 12.**
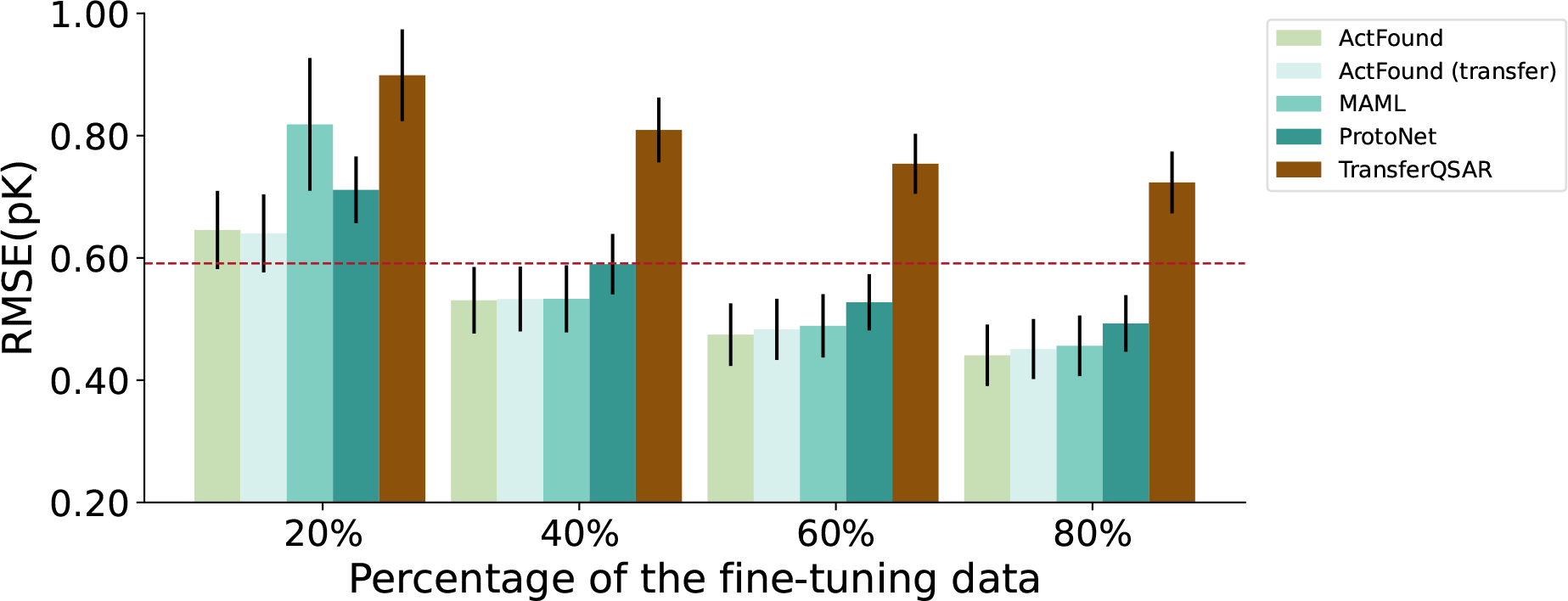
Bar plots comparing the bioactivity prediction on FEP benchmarks in terms of RMSE when 20%, 40%, 60%, 80% of the experimental bioactivity is used for fine-tuning. All models are trained on BindingDB. The red dashed line represents the computational results of FEP+(OPLS4).

**Supplementary Figure 13.**
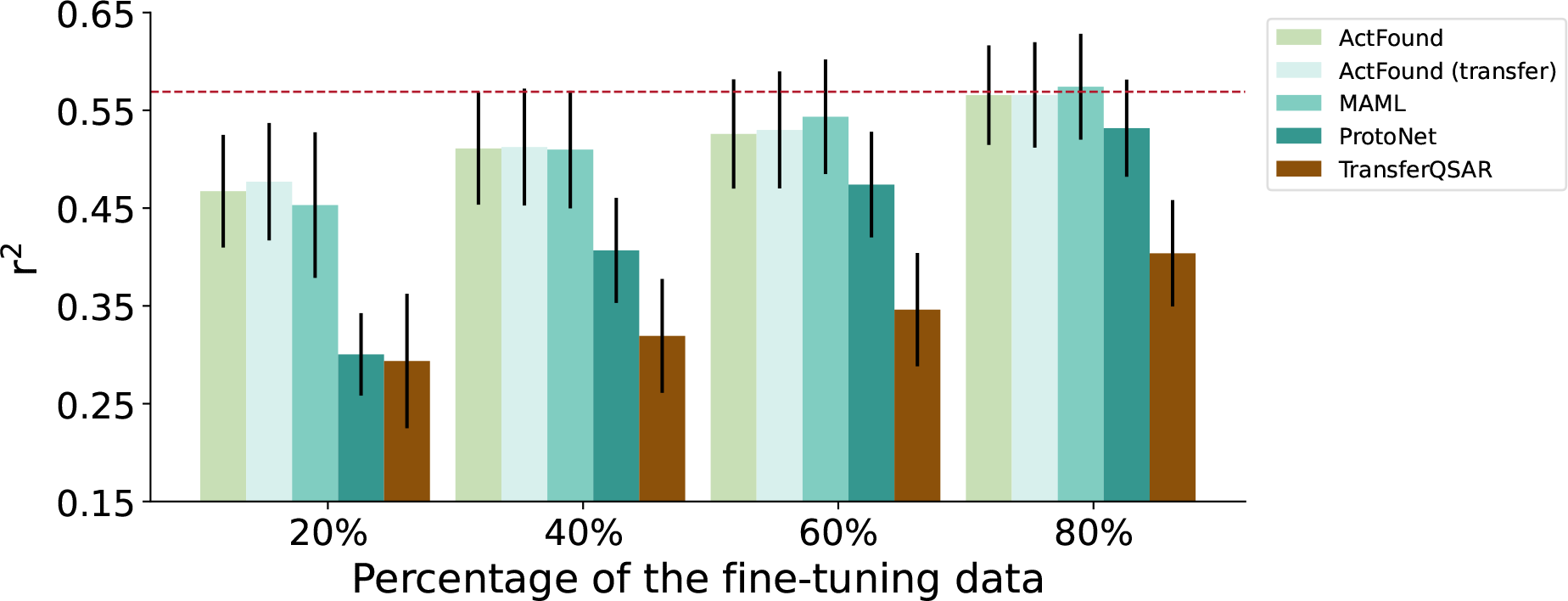
Bar plots comparing the bioactivity prediction on FEP benchmarks in terms of *r*^2^ when the FEP+(OPLS4) calculation results are used for 20%, 40%, 60%, 80% of the finetuning set. All models are trained on BindingDB. The red dashed line represents the computational results of FEP+(OPLS4).

**Supplementary Figure 14.**
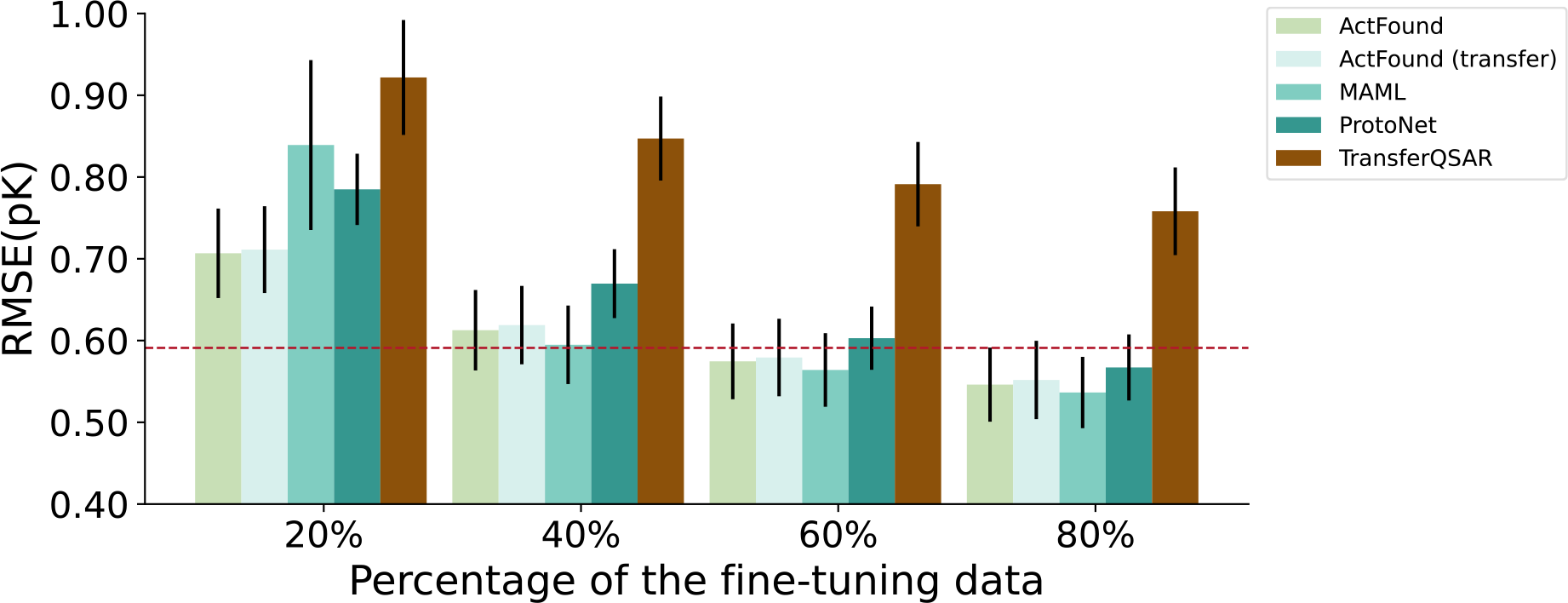
Bar plots comparing the bioactivity prediction on FEP benchmarks in terms of RMSE when the FEP+(OPLS4) calculation results are used for 20%, 40%, 60%, 80% of the fine-tuning set. All models are trained on BindingDB. The red dashed line represents the computational results of FEP+(OPLS4).

